# High-resolution analysis of Merkel Cell Polyomavirus in Merkel Cell Carcinoma reveals distinct integration patterns and suggests NHEJ and MMBIR as underlying mechanisms

**DOI:** 10.1101/2020.04.23.057703

**Authors:** Manja Czech-Sioli, Thomas Günther, Marlin Therre, Michael Spohn, Daniela Indenbirken, Juliane Theiss, Sabine Riethdorf, Minyue Qi, Malik Alawi, Corinna Wülbeck, Irene Fernandez-Cuesta, Franziska Esmek, Jürgen C. Becker, Adam Grundhoff, Nicole Fischer

**Affiliations:** Institute of Medical Microbiology, Virology and Hygiene, University Medical Center Hamburg-Eppendorf, Hamburg, 20246, Germany; Heinrich Pette Institute, Leibniz Institute for Experimental Virology, Hamburg, 20251, Germany; Institute of Tumorbiology, University Medical Center Hamburg-Eppendorf, Hamburg, 20246, Germany; Bioinformatics Core, University Medical Center Hamburg-Eppendorf, Hamburg, 20246, Germany; Translational skin cancer research, German Cancer Consortium (DKTK), University Hospital Essen, Essen, 45117, Germany; Deutsches Krebsforschungszentrum, Heidelberg, Germany; Institute of Nanostructure- and Solid State Physics (INF), Center for Hybrid Nanostructures (CHyN), University of Hamburg, Hamburg, 22761, Germany

## Abstract

Merkel Cell Polyomavirus (MCPyV) is the etiological agent of the majority of Merkel Cell Carcinomas (MCC). MCPyV positive MCCs harbor integrated, defective viral genomes that constitutively express viral oncogenes. Which molecular mechanisms promote viral integration, if distinct integration patterns exist, and if integration occurs preferentially at loci with specific chromatin states is unknown.

We here combined short and long-read (nanopore) next-generation sequencing and present the first high-resolution analysis of integration site structure in MCC cell lines as well as primary tumor material. We find two main types of integration site structure: Linear patterns with chromosomal breakpoints that map closely together, and complex integration loci that exhibit local amplification of genomic sequences flanking the viral DNA. Sequence analysis suggests that linear patterns are produced during viral replication by integration of defective/linear genomes into host DNA double strand breaks via non-homologous end joining, NHEJ. In contrast, our data strongly suggest that complex integration patterns are mediated by microhomology-mediated break-induced replication, MMBIR.

Furthermore, we show by ChIP-Seq and RNA-Seq analysis that MCPyV preferably integrates in open chromatin and provide evidence that viral oncogene expression is driven by the viral promoter region, rather than transcription from juxtaposed host promoters. Taken together, our data explain the characteristics of MCPyV integration and may also provide a model for integration of other oncogenic DNA viruses such as papillomaviruses.

**Author summary:** Integration of viral DNA into the host genome is a key event in the pathogenesis of many virus-induced cancers. One such cancer is Merkel cell carcinoma (MCC), a highly malignant tumor that harbors monoclonally integrated and replication-defective Merkel cell polyomavirus (MCPyV) genomes. Although MCPyV integration sites have been analyzed before, there is very little knowledge of the mechanisms that lead to mutagenesis and integration of viral genomes. We used multiple sequencing technologies and interrogation of chromatin states to perform a comprehensive characterization of MCPyV integration loci. This analysis allowed us to deduce the events that likely precede viral integration. We provide evidence that the mutations which result in the replication defective phenotype are acquired prior to integration and propose that the cellular DNA repair pathways non-homologous end joining (NHEJ) and microhomology-mediated break-induced replication (MMBIR) produce two principal MCPyV integration patterns (simple and complex, respectively). We show that, although MCPyV integrates predominantly in open chromatin regions, viral oncogene expression is independent of host promoters and driven by the viral promotor region. Our findings are important since they can explain the mechanisms of MCPyV integration. Furthermore, our model may also apply to papillomaviruses, another clinically important family of oncogenic DNA viruses.

## Introduction

Merkel cell carcinoma (MCC) is a rare but highly aggressive skin cancer occurring predominantly in elderly and immunosuppressed patients. The tumor shows a high propensity to metastasize, which is reflected in a poor 5 year survival rate [1-5]. The majority of MCCs (∼60-80% in the northern hemisphere) are causally linked to infection with Merkel cell polyomavirus (MCPyV). This notion is supported by the following observations: (i) all tumor cells in MCC harbor monoclonally integrated viral genomes [3, 6, 7], (ii) these integrated viral genomes carry tumor-specific mutations (see below) that are not present in viral episomes recovered from healthy individuals [3, 8, 9] and (iii) tumor cell viability is strictly dependent upon constitutive expression of viral oncoproteins from integrated genomes [10]. Interestingly, virus positive MCCs (VP-MCCs), in contrast to virus negative MCCs (VN-MCCs) show a rather low mutational burden in the host genome and lack typical cancer-driving alterations, indicating that viral oncoprotein expression in VP-MCCs is not only necessary but also largely sufficient for tumorigenesis [11-14].

In a healthy, immunocompetent person, the virus persists in an episomal form in a so-far unknown reservoir that most likely is located in the skin [15, 16]. Reactivation of such pools in immunosuppressed individuals is thought to favor the mutagenesis and genomic integration of viral DNA, two (presumably rare and independent) events that represent a prerequisite for MCC pathogenesis. The MCC-specific mutations present as point mutations and indels within the early region of the integrated viral genome and unequivocally result in the expression of a truncated Large T (LT) protein, LTtrunc [8, 17]. These truncated LT proteins are unable to support replication of viral DNA but preserve the ability to inactivate the tumor suppressor Retinoblastoma protein (pRb) via an amino terminal LxCxE motif. Integration sites vary between individual tumors and thus do not directly contribute to transformation. Furthermore, no clear integration, hot spots or regions have been identified [2, 18-21]. The transforming potential of the viral tumor antigens small T Antigen (sT) and LTtrunc have been the focus of a variety of studies [1, 22-24]. However, the mechanisms contributing to tumor-specific viral early gene mutation, as well as those that lead to viral integration are unclear. Likewise, it is unknown whether inactivating mutations occur before or after the integration of viral genomes. Viral DNA integration is a key event in several DNA tumor virus-associated cancers such as HPV-associated cervical cancer and HBV associated hepatocellular carcinoma [8, 9, 11, 25-27]. In both cancer types, viral integration sites are randomly distributed throughout the genome and integration is associated with deregulated viral oncogene expression. In HBV-induced hepatocarcinoma, aberrant expression of the viral HBx gene contributes to transformation. Similar to LTtrunc expression in MCC, in HPV-associated tumors oncoprotein E6/E7 expression is typically deregulated through a loss of E2 expression in HPV induced malignancies. Using Fiber FISH technology, viral integration with focal amplification of flanking host regions was recently demonstrated for the cell line 20861 (a subclone of the W12 cell line containing integrated HPV16 DNA) [28]. The integration locus in these cells was furthermore shown to exhibit epigenetic changes, resulting in the formation of super-enhancer elements which drive transcription of the viral oncoproteins.

Previous studies in MCC cell lines or primary material have used either DIPS-PCR, a ligation-mediated PCR assay [18-20], or short-read second generation sequencing [21, 29, 30] to detect MCPyV integration sites. While DIPS-PCR is rather labor-intensive and often fails to recover all breakpoints or resolve the structure of integration sites, the recent development of hybrid capture probe enrichment combined with short-read sequencing (capture sequencing) allows robust localization of viral integration sites [30].

In this study, we have characterized the integration pattern of 11 MCC cell lines and one primary tumor and its metastasis by second generation capture sequencing, third-generation nanopore sequencing, and the recently developed nanochannel sequencing technique. We show that MCPyV integration events can be assigned to one of two general groups based on their genomic arrangement: The first group is characterized by relatively simple, linear integration patterns presenting as single or concatemeric viral genomes flanked by host junctions that are positioned in close proximity two one another, suggesting that integration did not result in a major loss or amplification of flanking host sequences. In the second class, flanking cellular DNA is amplified, thus leading to more complex integration patterns in which the virus-host junctions are thousands of base pairs apart from each other. We provide evidence that host genomic amplifications in the latter group result from microhomology-mediated break-induced replication (MMBIR) [31] and propose a model that explains the different integration patterns of MCPyV in MCC.

Furthermore, we have performed ChIP-Seq analysis of histone modifications in the prototypic MCC cell lines MKL-1 and WaGa and show that the epigenomes of these two cell lines are highly similar. By cross-comparing the MCC histone modification landscape with ENCODE epigenome data across all identified integration loci, we provide evidence that MCPyV integration predominantly occurs in open chromatin that is devoid of constitutive heterochromatin marks. Our data furthermore suggest that the integration does not significantly alter the epigenetic landscape of flanking host loci, and that transcription from viral promoter elements is responsible for constitutive oncoprotein expression in MCC cells.

## Results

### Capture sequencing analysis of viral mutations and polymorphisms in MCC cell lines and primary tumor material

To allow high-resolution analysis of MCPyV integrates, we first enriched viral sequences and flanking host fragments from VP-MCC cell lines by hybridization capture and subsequent Illumina short-read sequencing (capture-sequencing). Hybridization capture was performed using 120mer SureSelect RNA capture probes, tiled along the entire MCPyV genome with a single nucleotide shift. We included four MCC cell lines with partially known integration sites identified by DIPS-PCR [20] (WaGa, LoKe, BroLi, PeTa) and seven cell lines with so far unknown viral and cellular breakpoints (MKL-1, MKL-2, WoWe-2, UKE-MCC-1a, UM-MCC-29, UKE-MCC-4a, UM-MCC-52). Furthermore, we included an MCC primary tumor and its bone metastasis (MCC-47T and -M, respectively). Coverage plots of viral reads aligned to the MCPyV genome (Genbank: JN707599) confirmed efficient enrichment and high read coverage (between 9,500 and 310,000) across all samples (Fig 1).

**Fig 1:**
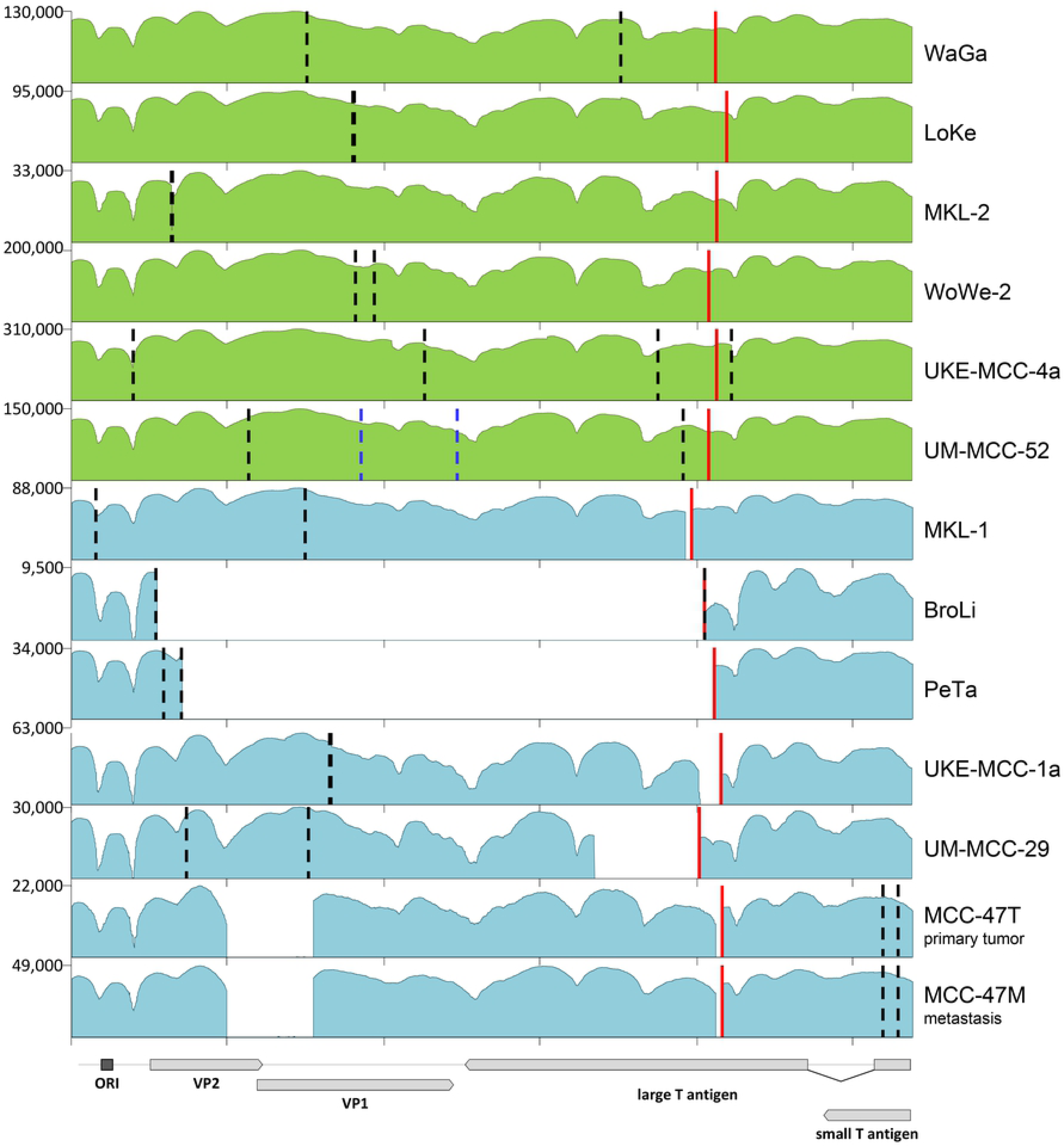
Coverage of capture sequencing reads to the MCPyV genome. Shown are log scale coverage plots of all viral reads aligned to the MCPyV genome (JN707599). The viral genome structure is illustrated in the lower panel. Red lines mark positions of LT truncations. Coverage plots shown in green represent samples with point mutations resulting in a stop codon and premature LT protein whereas coverage plots in blue show samples in which deletions or inversions cause frameshifts and subsequent premature stop codons. Dashed lines in black indicate breakpoints of the MCPyV genome into the host genome. In UKE-MCC-4a four breakpoints into Chr20 were detected. In UM-MCC-52 dashed lines mark breakpoints into Chr5 (blue) and Chr4 (black).

Variant calling readily identified sample-specific mutations, including those that lead to MCC-specific truncation of the large T open reading frame (marked by a red line in Fig 1; see Table 1 for the exact LT truncating events in each sample). The seven samples shown in green harbor point mutations that create a premature stop codon, whereas those shown in blue exhibit deletions (in case of PeTa and MCC-47 combined with inversions) that result in frameshifts and subsequent LT truncation. As expected, all truncating mutations preserve the LxCxE motif but remove the carboxyterminal origin-binding domain and helicase domains [32].

**Table 1:**
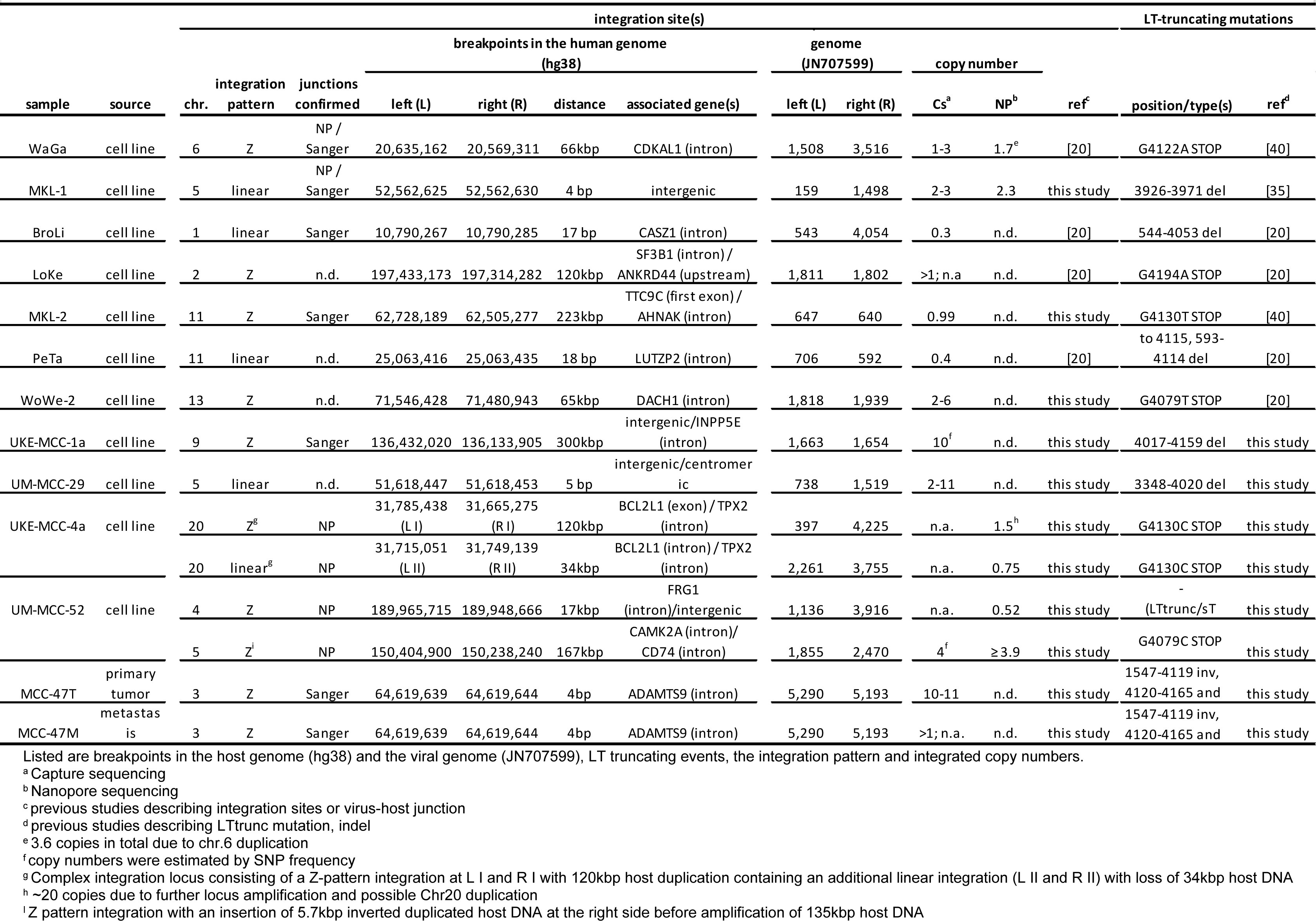
Integration sites of MCPyV in MCC samples.

The substantial read coverage levels achieved by capture sequencing additionally allowed us to perform high-confidence variant calling to evaluate potential viral genome heterogeneity within samples (see S1 Table for a complete list of variants). According to this analysis, WaGa, BroLi, MKL-2, UM-MCC-29, and UKE-MCC-4a each harbor distinct variant signatures with frequencies >99%, indicative of all viral copies within each sample being identical. Similarly, we find identical integrated viral genomes in the primary tumor MCC-47T and its descendent metastasis, MCC-47M. Interestingly, in three cell lines, we find additional lower frequency variants: A duplication in UKE-MCC-1a (bp 1,372-1,398 at 9.2% frequency), three point mutations in UM-MCC-52 (positions 1,708, 1,792 and 1,816 with frequencies of 15.8%, 20.2% and 22.1%, respectively), four point mutations in WoWe-2 (positions 3,784, 3,791, 3,812, 3,827 with frequencies of 74,3%, 74,8%, 80,4% and 80,5% respectively) and a deletion in UKE-MCC-4a (bp 2,053-3,047 at ∼33% frequency). These variants are not detected in any other sample, making contamination unlikely and suggesting integration of different MCPyV variants, or diversification by mutations occurring after the integration event.

### Analysis of virus-host breakpoints by short-read sequencing

To pinpoint MCPyV integration sites we mapped virus-host fusion reads from capture sequencing of the different MCC cell lines and the tumor and its metastasis to the human genome. In Fig 2A, we present an overview of integration sites across the human genome. Table 1 lists precise nucleotide positions and associated viral breakpoints, as well as genomic features at integration sites. S1 Fig shows the sequences of all identified virus-host junctions. Most integration sites are found in introns, some in intergenic or centromeric regions and one maps to an exon. Overall, we did not observe obvious overrepresentation of distinct genomic loci among integration sites (Fig 2A), a finding which is in accordance with previous studies [3, 18-20, 30, 33, 34]. Interestingly, however, we find that three cell lines harbor viral integrates in Chr5. While the number of samples investigated here is too small to allow calculation of statistical significance, we note that a recent study investigating a large cohort of VP-MCCs had suggested that Chr5 might be more prone to MCPyV integrations [30], a notion which seems to be supported by our observations.

**Fig 2:**
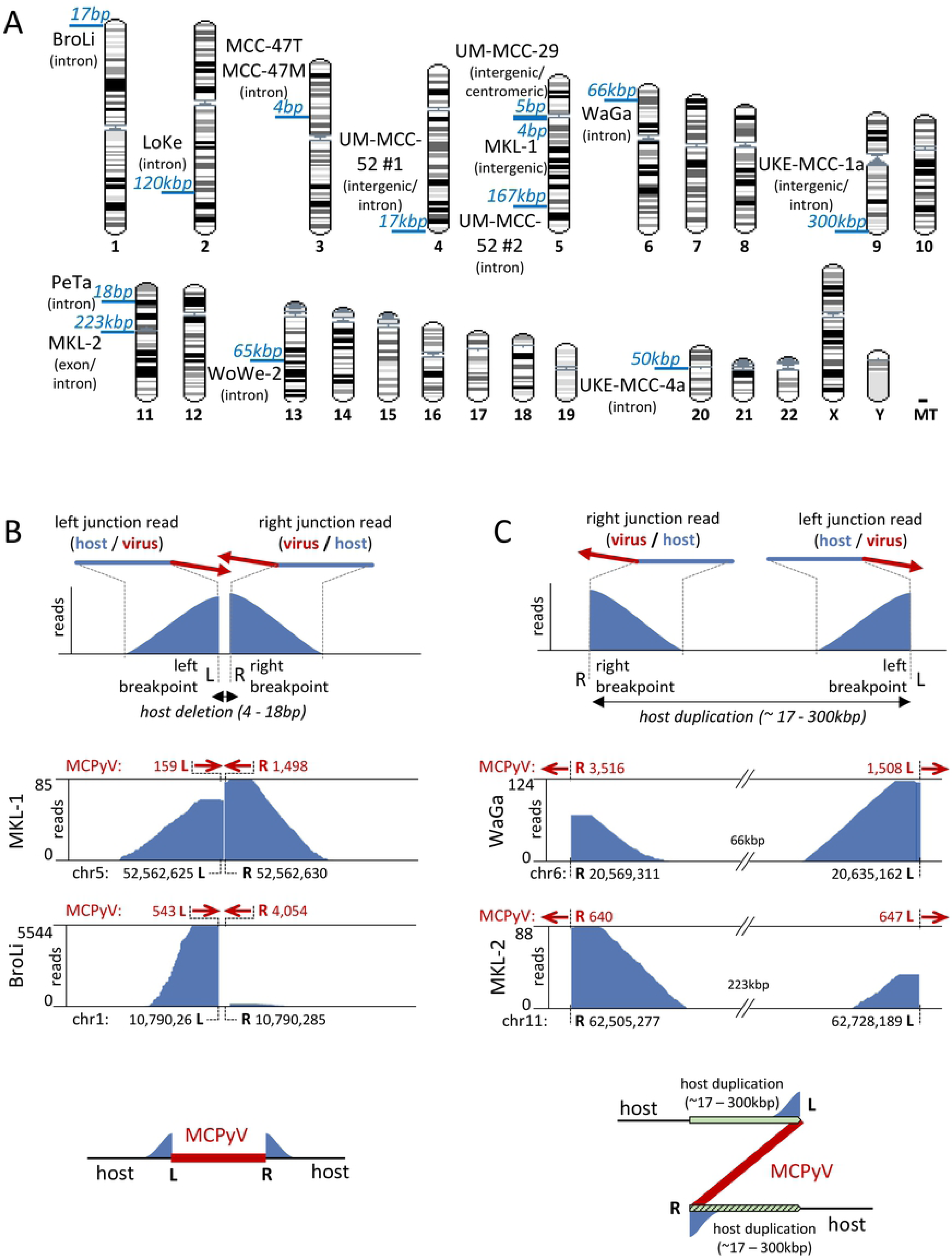
MCPyV integration sites detected by capture sequencing. (A): MCPyV integration sites in the human chromosomes. Depicted in blue is the distance between breakpoints on the host genome. Characteristics of the host genome at the breakpoints are indicated in brackets. (B) and (C): Schematic representation of the two characteristic groups of coverage profiles obtained by the mapping of virus-host fusion reads to the human genome. A schematic of virus-host fusion reads is depicted above the coverage patterns; red arrows indicate the direction of the MCPyV sequence in the fusion reads. (B) represents the first group characterised by short distances (4-18bp) between breakpoints on the host genome and inward-facing orientation of the fused viral sequences (upper panel). The middle panel shows coverage tracks from the cell lines MKL-1 and BroLi as examples. The bottom panel depicts a schematic model of the linear integration pattern deduced from the coverage profiles presented above. (C) represents the second group where host sequences in fusion reads map with large distances (17kbp to 300kbp) on the host genome and viral sequences show an outward-facing orientation. WaGa and MKL-2 coverage tracks are shown as examples with a schematic model of the integration pattern deduced from the coverage profiles above. Large host regions preceding the left virus-host junction are duplicated after the right virus-host breakpoint leading to a “Z” shape of the integration. Coverage tracks from all additional samples are provided in S4 Fig.

We unambiguously identified a single integration locus with two virus-host junctions for the majority of MCC cell lines (WaGa, MKL-1, BroLi, LoKe, MKL-2, PeTa, WoWe-2, UKE-MCC-1a, and UM-MCC-29). The primary tumor and its descended metastasis (MCC-47) also show a single integration locus that is identical between both samples. Based on nucleotide insertions at the virus-host junctions, we can distinguish three junction types. (i) junctions with no additional nucleotide insertions (MKL-2, PeTa, UM-MCC-29, BroLi, WoWe-2 and MCC-47T/M); (ii) 2-3 bp insertions of heterologous origin at one of the junctions (UKE-MCC-1a and LoKe) and (iii) 10-30 bp insertions of host (WaGa, originating from the second junction) or viral (MKL-1 and UKE-MCC-1a) sequences that are found immediately at or close to one of the junctions (see S1 Fig for more details).

### UM-MCC-52 and UKE-MCC-4a represent MCC cell lines with multiple integration sites

We found four virus-host junctions in UM-MCC-52 and UKE-MCC-4a, indicating the presence of two integration sites. In UM-MCC-52, the two sites mapped to Chr4 and Chr5, with 1bp insertions at the junctions on both chromosomes (Figs 1, 2A, S1 Fig, Table 1). While our variant analysis suggested that a fraction (∼19.4%) of viral genomes harbors three SNPs in VP1 (positions 1,708, 1,792 and 1,816), the SNPs were not present in any of the junction reads and capture sequencing therefore did not allow us to assign them to one of the loci (S2 Fig). In the cell line UKE-MCC-4a, all junction reads mapped to a 120kbp locus on Chr20 (Figs 1, 2A, S1 Fig and Table 1), again with heterologous nucleotide insertions at the virus-host junctions (see S1 Fig for more detailed information). Mutations in the early viral region were present in all reads while only ∼30% of the reads contain a deletion at position 2053 to 3047 of the viral genome. Interestingly, the viral breakpoint of one of the virus-host junctions is located within this deletion, indicating it must be absent from the viral copy located at this junction (S3 Fig).

### Analysis of integration patterns by capture sequencing

As shown in Figs 2B and -C, mapping of virus-host fusion reads to the reference human genome produced two distinct alignment patterns, each with a characteristic coverage profile. The upper panels in Figs 2B and -C show a schematic depiction of each coverage pattern, while representative data from two cell lines belonging to each group (see S4 Fig for the residual samples belonging to the two groups of coverage patterns) are shown underneath. A schematic depiction of the deduced integration site structure is shown in the bottom panels.

In the first group, reads spanning the breakpoints mapped closely together (4-18 bp distance), with inward-facing orientation of the fused viral sequences. The associated host coverage profiles present as a split peak with a central gap that separates the breakpoints (see MKL-1 and BroLi as examples in the center panels of Fig 2B). This pattern is suggestive of a linear viral integration event in which a few bases of host DNA have been lost (Fig 2B, bottom panel). Similar patterns were identified for PeTa and UM-MCC-29 cell lines (S4A Fig).

The second group is characterized by breakpoint-spanning reads that typically map in much greater distance from one another (17kbp to 300kbp), with viral sequences that extend in an outward-facing orientation (Fig 2C, top). The simplest explanation for such a coverage pattern is a duplication of the host DNA between the two breakpoints, leading to an integration pattern resembling a “Z” shape (Fig 2C, bottom panel). Hence, reads originating from the left and right junctions of the integration site align with the reference genome in a seemingly inverted manner, with the right junction reads preceding those from the left junction. In addition to the WaGa and MKL-2 samples shown in the center panels of Fig 2C, the samples LoKe, WoWe-2, UKE-MCC-1a and MCC-47T and -M (tumor and metastasis) also show a Z-pattern integration (S4B Fig). The MCC-47 samples differ from the others in that the duplicated host sequence is only 6bp in length (S1, S4C, S5 Figs), indicating that duplication of tens or hundreds of kbp is a frequent, but not a generally valid feature of this type of integration pattern.

### Calculation of MCPyV genome copy numbers based on capture sequencing data

Previous studies reported integration of multiple copies of MCPyV genomes arranged as concatemers [20, 30, 34, 35]. We estimated the number of integrated MCPyV copies data by calculating the number of virus-host junction reads relative to viral reads which encompass breakpoint positions, but do not contain host junction sequences (referred to as fusion or virus-only reads, respectively, in the following). In cells harboring a single integration locus virus-only reads must necessarily be derived from internal virus copies, and the read count ratio thus can provide an estimate of concatemeric unit numbers.

In BroLi and MKL-2 we only find fusion reads at breakpoints, indicating integration of a single (partial) viral copy. In the case of BroLi this integrate lacks two-thirds of the viral genome, whereas in MKL-2 only 6bp are missing. In contrast, we find high numbers of virus-only reads covering the breakpoints in WaGa, MKL-1, LoKe, WoWe-2, UKE-MCC-1a, UM-MCC-29, MCC-47T and MCC-47M, suggesting the presence of viral concatemers. Estimated copy numbers of viral genomes in integrated concatemers are listed in Table 1 and range between two and 11 copies. For LoKe and UKE-MCC-1a copy numbers could not be estimated due to breakpoints in the viral genome being too close to one another. Interestingly, variant calling revealed that in UKE-MCC-1a, a duplication at viral position 1373-1398 is only present in 9.2% of the reads, suggesting that only a fraction of the viral copies contains the duplication. Sanger sequencing of an 800bp PCR product covering the virus-host junction revealed that the copy closest to the left junction contains the 25bp duplication. This duplicated sequence is also inserted directly at the junction (S1 Fig) suggesting that the duplication inside the viral genome was acquired during the integration process, only in the viral copy closest to the virus-host junction. Based on the frequency of the duplication (9.2%) we estimate an integration of 10 viral genomes in the case of UKE-MCC-1a. The samples UKE-MCC-4a and UM-MCC-52 contain multiple integration sites. Thus, we were unable to distinguish between virus-only reads derived from internal concatemer copies or the other integration sites.

### Analysis of potential integration locus amplification

Studies for papillomavirus integration in cervical cancer showed that entire integration loci and flanking host DNA can be amplified several times [26, 28, 36]. Since FISH analysis for the MCPyV genome consistently yielded two signals in WaGa cells compared to one signal in MKL-1 cells [24, 37] we investigated if the complete integration locus is amplified in WaGa cells. To calculate genomic host copy number variations, we used input data from ChIP-Seq analysis performed in WaGa and MKL-1 cells (see below), which resemble low coverage whole-genome sequencing data (WGS). Genome copy calculation reveals amplification of the entire Chr6 (including MCPyV integration) in WaGa cells (Fig 3A). Analysis of relative genomic copy numbers 60kbp up- and downstream of the integration locus (i.e. regions not affected by the duplication) suggests the presence of three copies of Chr6 (Fig 3B, left panel). Since the duplicated regions are found in five copies, this implies that it is the Chr6 copy carrying MCPyV which is duplicated. The two signals in WaGa cells observed by FISH analysis therefore represent Chr6 duplication rather than a specific amplification restricted to the integration locus. Although genomic amplifications are observed in some chromosomes in MKL-1 cells, we do not detect amplification of the entire Chr5, which carries the MCPyV integration site in this cell line (Fig 3A), or an amplification of host regions directly flanking the integration (Fig 3B, right panel). Additionally, we calculated integrated viral genome copy numbers for WaGa and MKL-1 from the ChIP-sequencing input data (Fig 3C), thereby confirming the results obtained by our estimation based on the capture sequencing data (WaGa 1-3 copies, MKL-1 2-3 copies, see also Table 1).

**Fig 3.**
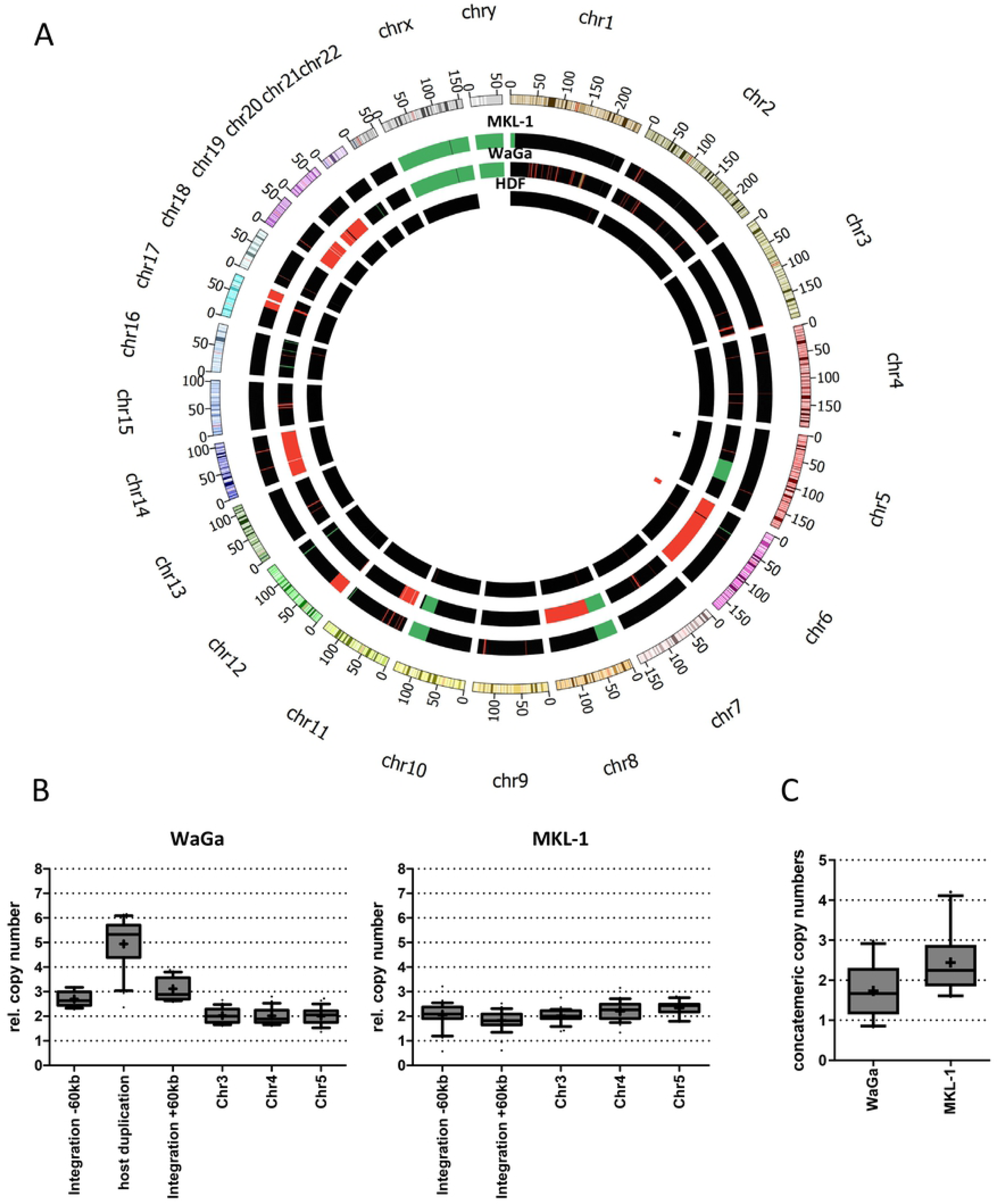
Viral copy number calculation in WaGa and MKL-1 cells. (A): Circos plot of copy number variations in WaGa and MKL-1 cells as calculated by FREEC using low coverage WGS data (ChIP-Seq input). The colour code indicates chromosome aberrations in fold haploid (black = 2n; green = 1n; red >= 3n; white = 0n). Female HDF cells are shown as control with n = 2. The position of MCPyV integrations are shown in the innermost circle (black: MKL-1; red: WaGa). (B): Normalized relative genomic DNA copy numbers immediately upstream (integration -60kbp) and downstream (+60kbp) of the respective MCPyV integration sites are shown in comparison to three indicated genomic control sites of the same length (Chr3, 4 and 5). Additionally, the 60kbp host duplication of WaGa cells is shown. Normalized data are presented as box and whisker plots of 5kb shifting windows (shift size = 2.5kbp) across the respective region of interest with median (horizontal line) and average (indicated by “+”). (C): Concatemeric copy numbers within each integration site in WaGa and MKL-1 were calculated from ChIP-Seq input data as described in the materials and methods section. Normalized data are shown as a box and whiskers plot of 1kbp shifting windows (shift size 0.5kbp) across the MCPyV reference genome (JN707599).

### Nanochannel and Nanopore sequencing confirm integrated copy numbers and reveal integration patterns of MCPyV in WaGa and MKL-1 cells

Short-read capture sequencing provides exact information on breakpoint location and viral sequence variants but is limited in terms of exact determination of integrated viral copy numbers and integration patterns. To confirm estimated copy numbers and linear or Z integration patterns as deduced from capture sequencing, we performed nanochannel and nanopore sequencing on a subset of samples. Nanochannel sequencing employs optical mapping of single DNA molecules and allows for fast determination of copy numbers and longitudinal sequence patterns within long DNA fragments [38]. The method is based on hybridization of fluorescently labelled probes, which are hybridized with high-molecular weight genomic DNA (HMW). The DNA is subsequently threaded through nanochannels, and detectors lining the channel are used to measure fluorescent signals along the length of intact DNA molecules. Nanochannel sequencing has kbp rather than bp resolution but does not require mechanic manipulation or amplification during library preparation, thus reducing the risk of introducing experimental artefacts. We analyzed HMW DNA from MKL-1 cells (a cell line with a linear integration pattern), which was fluorescently labelled with two LT-specific ATTO647N-probes. Fig 4A shows the measurement of a roughly 90kbp DNA fragment with a specific fluorescence peak detected over a period of 1.8ms, which corresponds to a size of approximately 17kbp. The measurement is in very good agreement with the 2-3 copy number estimation calculated from capture sequencing and ChIP-sequencing input data, thus confirming these data via a completely independent method. We additionally subjected HMW DNA from MKL-1 cells to Oxford Nanopore sequencing. As shown in Fig 4B (upper panel), we obtained several reads mapping to the integration site, including a single 104kbp read, which spans the entire MKL-1 integration locus and flanking sequences. Analysis of this read shows an integrated concatemer with two complete (2x 5.4kbp) and one partial (4.1kbp) copies, thereby confirming the results obtained by nanochannel sequencing as well as the linear integration and the junction sequences as determined by capture sequencing.

**Fig 4.**
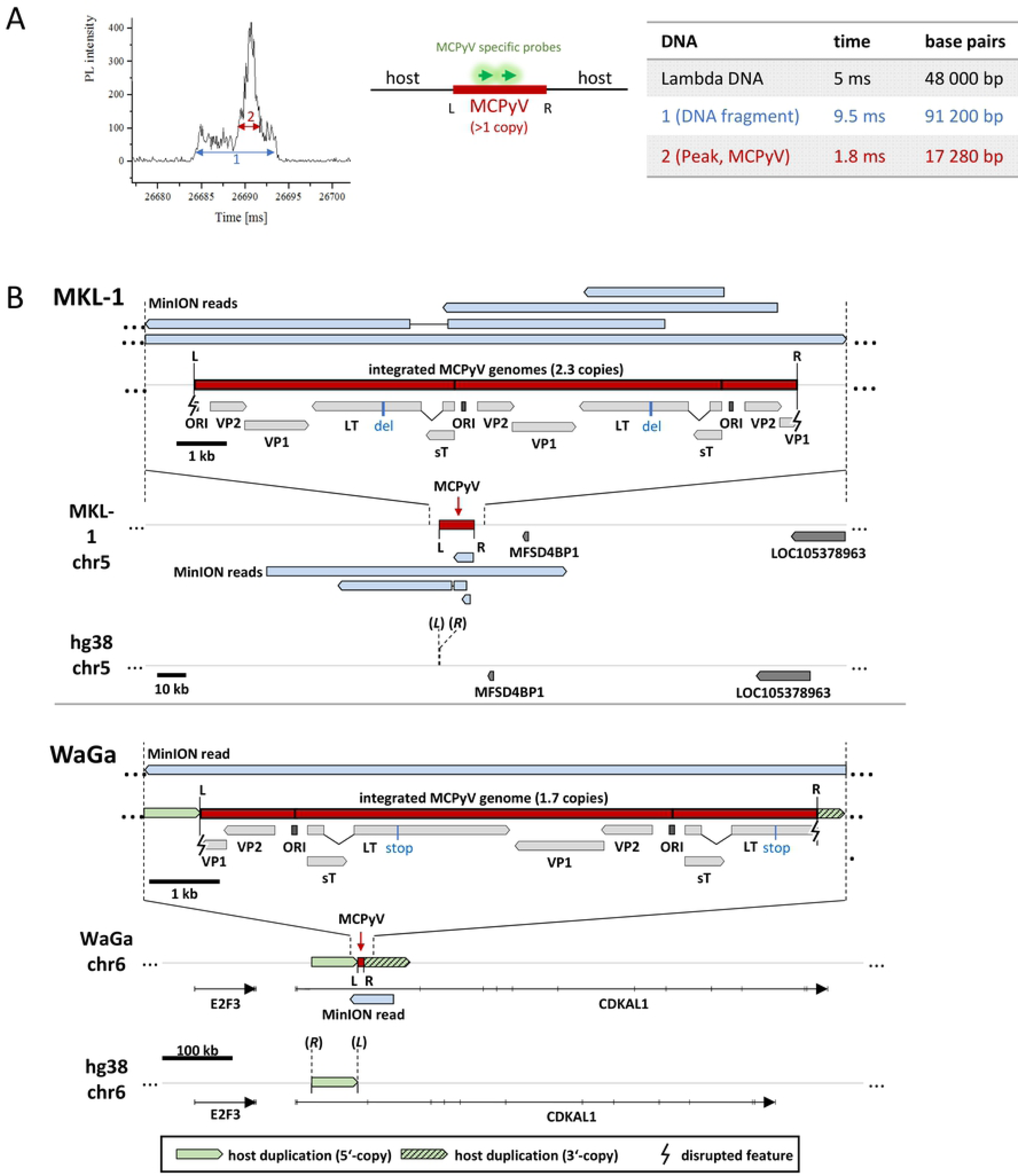
Nanochannel and nanopore sequencing determine viral integration patterns and copy numbers. (A): Optical signature (“barcode”) of a DNA fragment from MKL-1 cells. Shown is the time dependent intensity of the photoluminescence (PL intensity) of a single DNA fragment (1, blue), with an additional ATTO647N fluorescence peak (2, red). The fragment has a length of ∼ 90kbp, calculated after calibration with λ-DNA (48kbp) as a standard. The peak of ATTO647N fluorescence has a length of ∼ 17 kbp, corresponding to three integrated MCPyV copies (two complete copies, 5.4kbp each, and one partial copy with 4.1kb length). (B): Reads from nanopore sequencing for MKL-1 (upper panel) and WaGa (lower panel) mapped to the integration site of each cell line with an overview of the genomic locus in the reference genome (bottom), the integration locus as observed in the cell line (middle) and a close up on the integrated viral genome (top). For MKL-1, one read (104kbp in size) and three shorter reads cover the integration site. The long read confirms the linear integration of three concatemeric MCPyV copies (two full and one partial). For WaGa one 62kbp read covers the integration site. The read confirms the integration of two concatemeric MCPyV copies (one full and one partial) and the Z-pattern integration with duplication of the host sequence at the integration site. L and R indicate the left and right virus-host junction while (*L*) and (*R*) mark the position of the left and right junction sites in the host reference genome according to Table 1.

We also performed nanopore sequencing on HMW DNA from WaGa cells as a representative of the proposed Z-pattern integration (Fig 4B, lower panel). A 62kbp read covering the integration locus confirmed the Z-pattern integration with a large duplication of the host DNA between the two junction sites and the integration of two concatemeric viral copies (one complete and one partial genome). Again, nanopore sequencing confirms the validity of copy number estimates calculated from short-read sequencing (1.7 copies as determined by nanopore sequencing vs. 1-3 capture sequencing copy number estimation).

### Nanopore sequencing uncovers complex integration patterns in the MCC cell lines UM-MCC-52 and UKE-MCC-4a

In the cell line UM-MCC-52, we identified two integration sites in chromosomes 4 and 5. Our capture sequencing data are suggestive of a Z-pattern integration with duplications of 17 and 167kbp, respectively (Fig 5A and -B). However, while the Chr4 site shows the typical read mapping pattern as depicted in Fig 2C (Fig 5A), fusion reads from both Chr5 junctions have viral sequences that extend in the same direction when mapped to the reference human genome (Fig 5B, upper panel). Since there is no indication for an inversion within the MCPyV genome itself, these data are suggestive of a partial inversion of the 167kbp host duplication downstream of the right (R) junction (Fig 5B, lower panel). To verify this hypothesis and determine integrated viral copy numbers, we performed nanopore sequencing on HMW DNA of UM-MCC-52 and detected several reads that cover the integration loci (Fig 5C and - D). As expected, Chr4 shows a Z-pattern integration with a 17kbp host duplication. Interestingly, this site harbors one partial MCPyV genome (bp 1,136 to 3,916) that only contains distal (3’) fragments of the LT and VP1 ORFs, whereas the entire NCCR and the proximal (5’) early region encoding LTtrunc and sT are missing (Fig 5C). This partial genome also contains the three SNPs at positions 1,708, 1,792 and 1,816 that had been identified in our capture sequencing analysis (S2 Fig). At the Chr5 site, we detected integration of a concatemer containing at least three complete and one partial MCPyV genomes (Fig 5D). As expected from short read sequencing, none of the viral genomes contains the three SNPs present at the integrated viral genome at Chr4 but all harbor identical LT truncating mutations. The SNP frequency (∼19%) at Chr4 supports the presence of four MCPyV copies in Chr5. Furthermore, the MinION reads confirm the suspected Z-pattern integration at Chr5 and show that indeed a 5.7kbp inverted duplicated host sequence originating from further upstream is inserted at the right virus-host junction followed by 135kbp of duplicated host DNA in direct orientation (Fig 5D). Taken together these results suggest that in UM-MCC-52 two (likely independent) integration events occurred on Chr4 and Chr5. The small fragment in Chr4 most likely does not contribute to transformation as it lacks the viral oncogenes.

**Fig 5.**
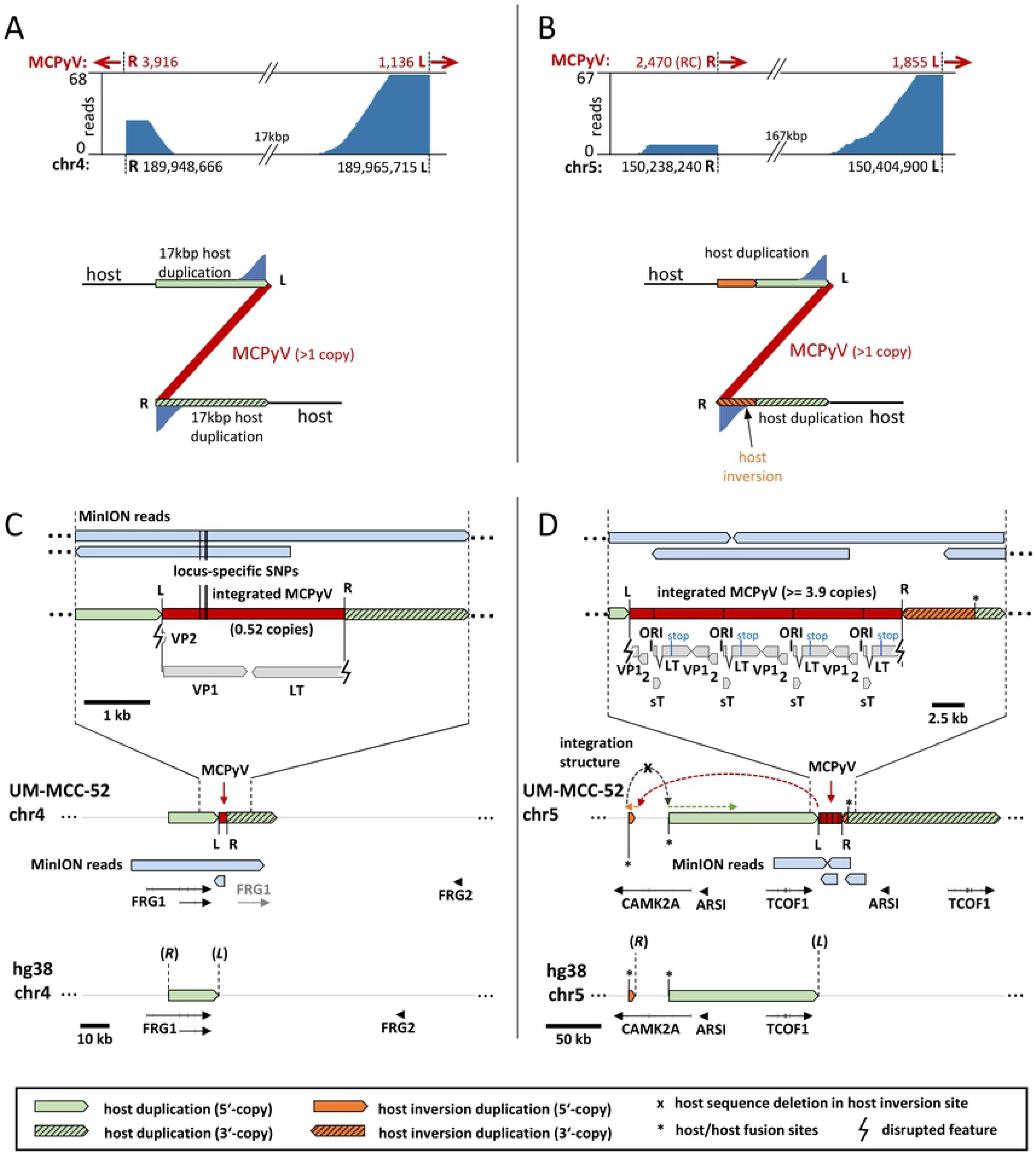
Complex integration pattern of UM-MCC-52. (A)+(B): MCPyV-host fusion reads from capture sequencing of sample UM-MCC-52 were mapped to the human genome. Shown is the coverage at the breakpoints in the host genome on Chr4 (A) and Chr5 (B). Red arrows indicate the direction of the viral sequences in the virus-host fusion reads. (RC)= Reverse complement orientation of MCPyV genome compared to the other junctions. Deduced integration patterns are shown below with a Z-pattern containing amplification of 17kbp host DNA in Chr4. The integration into Chr5 in addition to a Z-pattern must contain further inversions based on the read directions. As there is no indication for an inversion in the MCPyV genome, parts of host DNA at the right junction (R) must be inverted. (C)+(D): Reads from nanopore sequencing of UM-MCC-52 are mapped to both integration sites (Chr4, (C) and Chr5, (D)). In Chr4 0.52 MCPyV copies with three specific SNPs (bp 1,708; 1,792; 1,816; not present at the Chr5 integration) are integrated as a Z-pattern with duplication of 17kbp host DNA. In Chr5, MCPyV is integrated as a concatemer of at least 3.9 copies. MinION reads proof a Z-pattern integration with an insertion of 5.7kbp inverted duplicated host sequence at the right side that originates from 38kbp upstream of the 135kbp host sequence that is duplicated afterwards. Dashed coloured arrows indicate the complex structure of the integration locus. Duplicated host transcripts are shown in grey. L and R indicate the left and right virus-host junction while (*L*) and (*R*) mark the position of the left and right junction sites in the host reference genome according to Table 1.

In the cell line UKE-MCC-4a, our capture sequencing had identified a very complex 120kbp integration locus with four virus-host junctions (Fig 6A). The read orientation at the outmost junctions (R I and L I) together with long distances between breakpoints (120kbp) suggests a Z-pattern integration between these two junctions (site I in the following). The integrated viral genome at the inner junctions (L II and R II) is in a reverse complement orientation compared to the junctions R I and L I and shows inward-facing orientation of the viral sequences. Since there is no indication for an inversion within the MCPyV genome, a second linear integration between L II and R II (site II in the following) seems likely. While we hypothesized that a Z-pattern integration followed by a second linear insertion may have occurred at this locus, we could not resolve its structure based on short read data alone. To determine the correct structure of the MCPyV integration locus we again used Nanopore sequencing (Fig 6B). We obtained several reads that clearly support a Z-pattern with the duplication of 120kbp host sequence at integration site I (Fig 6B). One and a half MCPyV genomes are integrated at this site, but only the first copy harbors the deletion of bp 2053 to 3047 we already identified in our short-read sequencing (S3 Fig). Inside the 120kbp host duplication 34kbp are deleted and one partial copy of MCPyV is inserted in a linear fashion at site II. As expected from capture sequencing, this genome does not contain the deletion observed at site I but shares its LT inactivating mutations. This suggests that sites I and II contain the same MCPyV variant, and that deletion in the first copy of MCPyV at site I was likely acquired after the integration event. Hence, in contrast to the two integration events in UM-MCC-52, in UKE-MCC-4a sites I and II seem to be result from a single integration event. Downstream of the 120kbp duplication follows another integration of MCPyV (site I’ as it is identical to site I) with a second amplification of host DNA leading to the observed order I - II - I’ (Fig 6B). The increased coverage at the integration locus compared to the host genome (Fig 6C) suggests additional amplification of the complete locus. From our MinION data, we calculate a total of 20 copies for the complete locus (Fig 6D). Of note, the repetitive element is I - II, so the integration locus starts with site I and ends with site I’ as only reads from the sites I and I’ continue further into the host genome over the breakpoints of site II (positions of L II and R II). Reads from site II that reach the positions of the junctions R I and L I always contain the site I or site I’ integration, respectively.

**Fig 6.**
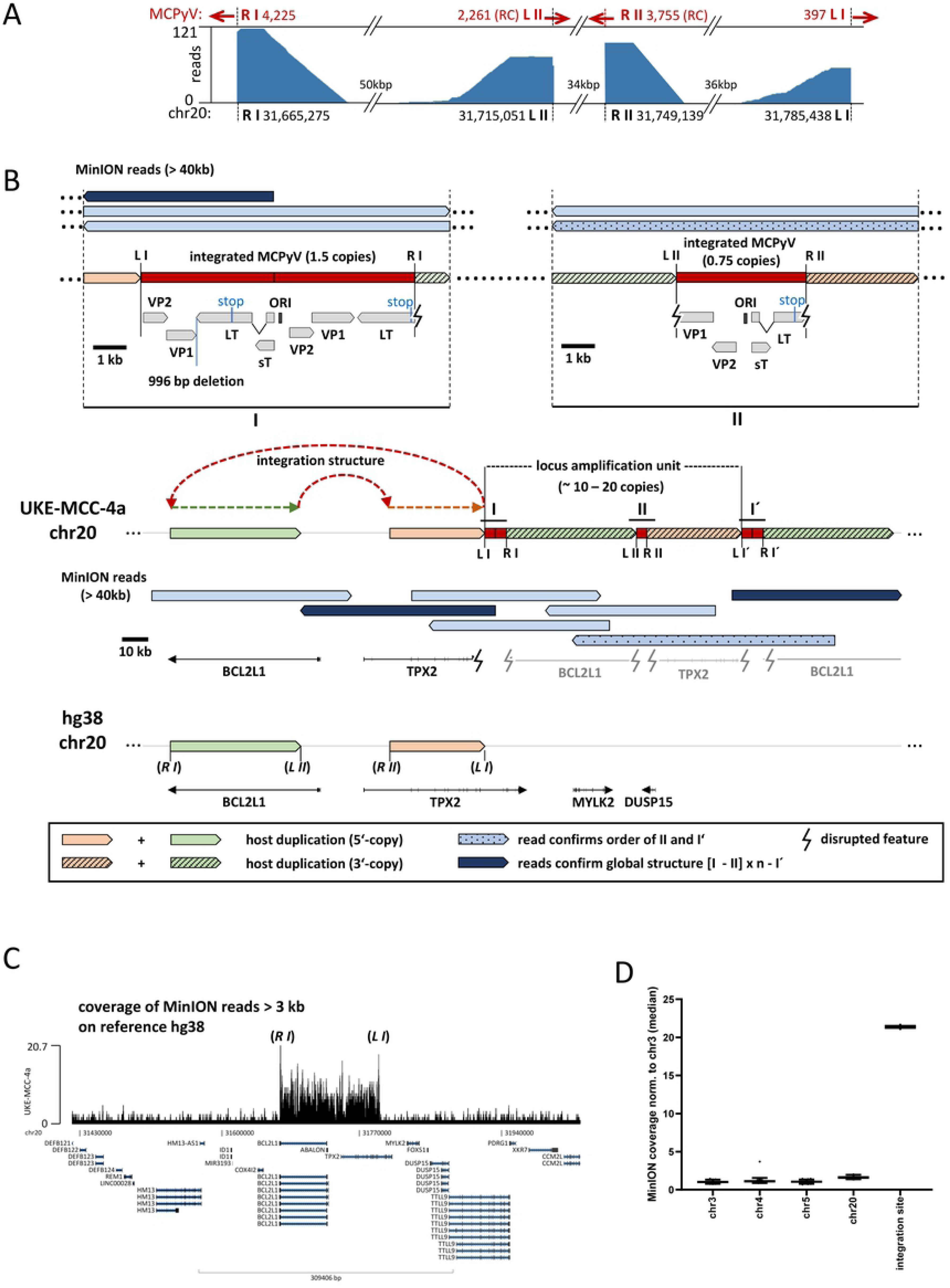
Complex integration pattern of UKE-MCC-4a. (A): MCPyV-host fusion reads from capture sequencing of sample UKE-MCC-4a were mapped to the human genome. Shown is the coverage at the four breakpoints in the host genome (R I, L II, R II and L I), red arrows indicate the direction of the viral sequences in the virus-host fusion reads. 81 Reads at junction R II are mapped by BLAST only (not by aligner). MCPyV reads that are reverse complementary (RC) fused to the host sequences (compared to the other breakpoints) are identified at L II and R II. (B): MinION reads >40kbp aligning to the integration site with an overview of the genomic locus in the reference genome (bottom), the integration locus as observed in UKE-MCC4a (middle) and a close up on the integrated viral genome at both integration sites (site I and site II) as confirmed by MinION reads (top). Site I shows a Z-pattern integration (amplification of 120kbp host DNA between R I and L I) of 1.5 concatemeric copies of MCPyV harboring a deletion of 996 bp only in the first of the two consecutive MCPyV copies. Site II shows a linear integration of 0.75 copies MCPyV (without the deletion) with a loss of 34kbp host DNA between L II and R II. The patterned read confirms the insertion of site II in the duplicated host DNA between R I and L I as well as a second insertion of site I (I’) with duplicated host DNA after the first Z-loop. The dark blue MinION reads confirm the order I – II – I’ since they continue from site I and site I’ into the host genome over the host positions of L II and R II of integration site II. The amplification unit is I – II (approximately 10-20 repeated units, see B and calculation in C). Dashed colored arrows highlight the structure of the complex integration product. Duplicated host features are shown in grey. L and R indicate the left and right sites of the virus-host junctions I and II while (*L*) and (*R*) mark the position of the left and right junction sites I and II in the host reference genome according to Table 1. (C): Coverage of MinION reads (with a size > 3 kbp) indicates amplification of the entire integration region. (D): Copy number calculation from MinION reads > 3 kbp in the integration region relative to multiple random regions on the indicated host chromosomes. Assuming a chromosome number of n=2 (most likely 3 for chr20) there may be either 10 large locus amplification units on both chromosomes of chr20 or 20 copies on only one chromosome of chr20.

### Microhomologies between viral and host sequences at integration sites

To understand the integration mechanism of MCPyV in more detail, we investigated the putative presence of microhomologies, as previously reported for papillomavirus integration sites [36, 39]. We therefore analyzed virus-host junctions from all integration sites in our study for matching bases between virus and host. While the occurrence of short homologies of 3bp directly at the junction of most samples is likely stochastic, we observed repetitive matching of short sequence stretches between virus and host that are intercepted by non-matching sequences of variable length (Fig 7A, S1 Fig). To detect and assess the matching sequences we developed a model that calculates homology scores for both sides of each virus-host junction, dependent on the distance from the junction. The sequences at the junction of viral and host origin are referred to as the virus side and the host side, respectively, in the following. Statistical analysis shows that in the case of Z-pattern integration, the virus side has significantly higher scores compared to scores obtained from 200 random sequences (Fig 7B). In contrast, the host side in the Z-pattern and viral and host sides in the linear integration pattern do not show significant homology compared to random sequences. These results suggest that in Z-pattern integration, microhomologies between viral and host sequences on the virus side of the resulting junction contribute to integration of MCPyV and that this initial integration step is different from linear integration.

**Fig 7.**
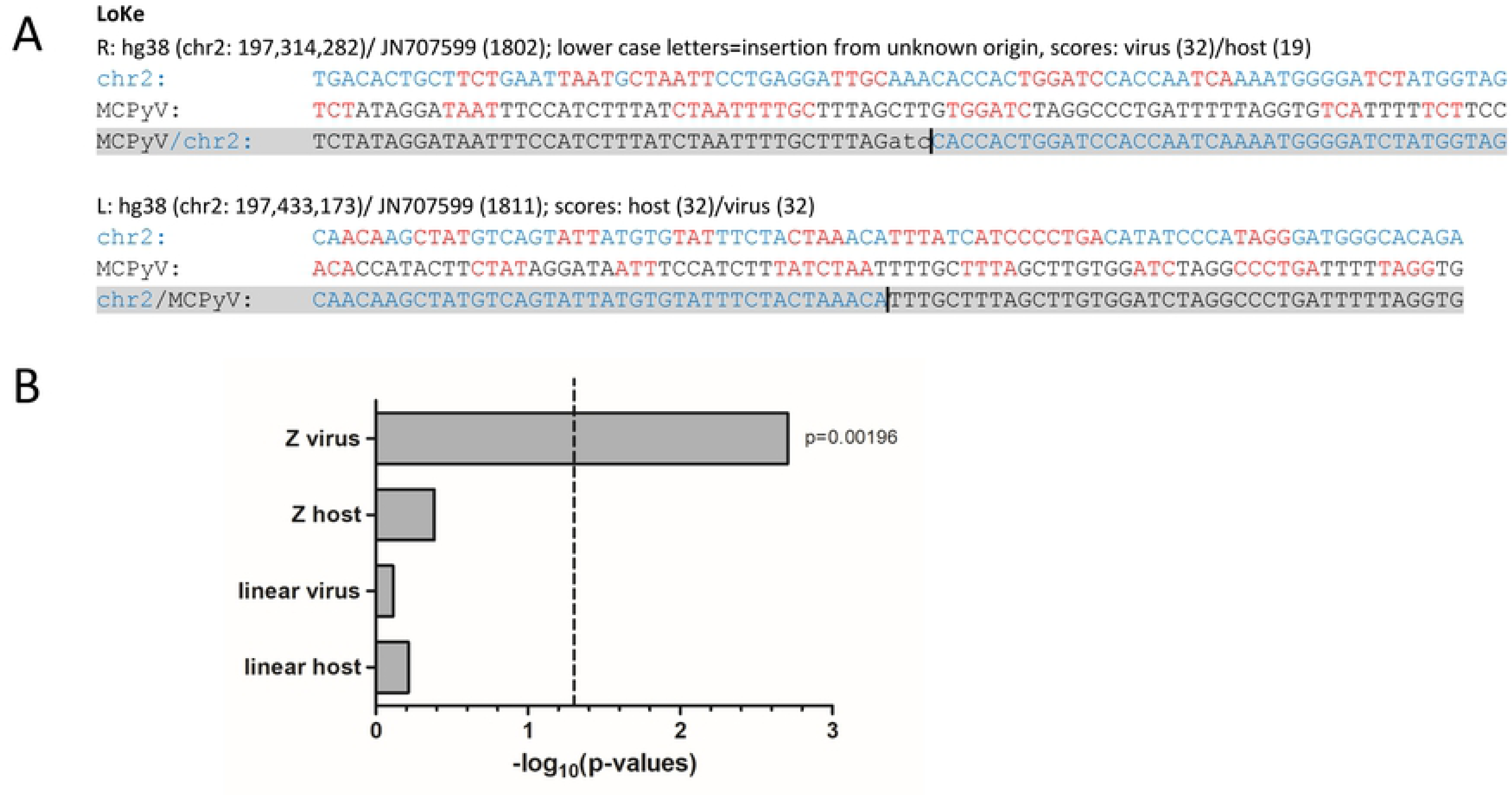
Microhomologies between virus and host sequences. (A): Virus-host junctions of the LoKe cell line. Sequences at the virus-host junction (in grey) were derived from capture sequencing and aligned to reference sequences for the human genome (hg38) and MCPyV (JN707599). Depicted are 40bp upstream and downstream from the virus-host junction (indicated by a black line, extended for 3bp at the right junction due to an insertion). Human sequences are shown in blue and viral sequences in black letters. Microhomologies are illustrated in red. Microhomology scores were calculated between the virus and host sequences for the virus side (viral sequence of the junction) and the host side (host sequence of the junction). All additional samples can be found in S1 Fig. (B): Scores from the virus and host side of samples showing Z-pattern or linear integration were compared to scores obtained for 200 random viral and host sequences. The virus side of Z-pattern integration shows significantly higher homology scores (p<0.05, dashed line). The host side and the linear integration pattern are not significantly different.

### Viral gene expression in MCC cell lines

Tumor cell proliferation in MCPyV positive MCCs is dependent on the constitutive expression of sT and truncated LT [3, 33, 35, 40, 41]. During viral replication, the T-antigens are expressed from the early viral promoter located in the non-coding control region (NCCR). Similar to previous results [30] we show that MCPyV integrates into diverse genomic regions (exons, introns, intergenic, centromeric) raising the question of whether the viral or an adjacent cellular promoter drives T-antigen expression. In addition, we sought to investigate whether the integration may perturb cellular gene expression at, or in close proximity to, the viral integration site. We therefore performed ChIP-Seq analysis of activating (H3K4-me3) and repressive (H3K27-me3) histone marks in WaGa and MKL-1 cells. WaGa cells harbor the viral integrate in the fourth intron of the gene CDKAL1, whereas in MKL-1 cells the viral genome is integrated in an intergenic region that does not harbor any annotated genes within a 300kbp distance. While we do not find H3K27-me3 to be present on integrated MCPyV, we find H3K4-me3 covering the entire viral early region in both cell lines (Fig 8A). This is clearly different from replicating viral genomes in PFSK-1 cells (Fig 8B) in which H3K4-me3 is present mainly on the NCCR and miRNA promoter region [37]. In contrast, in WaGa and MKL-1 cells the H3K4-me3 signals start at the early promoter and reach a plateau downstream of the LT/sT start codon, without the distinct enrichment observed at the miRNA promoter of actively replicating episomes. To investigate if early viral gene expression is driven by viral or cellular promoter elements we additionally performed transcriptome analysis of WaGa and MKL-1 cells. For this purpose, we mapped RNA-Seq reads to a reconstituted reference genome containing the identified integration sites and analyzed splices connecting to the splice acceptor of the second LT exon (S6 Fig). MKL-1 cells only showed canonical splice junctions between the first and second exons of LT (S6A Fig), a result which was expected due to the large distance between the integration site and the closest annotated host gene. In the case of WaGa we indeed detected some fusion reads between the second exon of LT and the splice donor of CDKAL1 exon4 (S6A Fig), but reads containing the canonical LT splice junction were 32 times more abundant. Together with the observed H3K4-me3 and RNA-Seq read coverage patterns (Fig 8A and S6B Fig, respectively), this suggests that the great majority of early MCPyV transcripts originate from viral promoter elements. Since the CDKAL1/LT fusion transcript furthermore is predicted to generate an out-of-frame product, expression of LTtrunc is likely to entirely depend on viral promoters.

**Fig 8.**
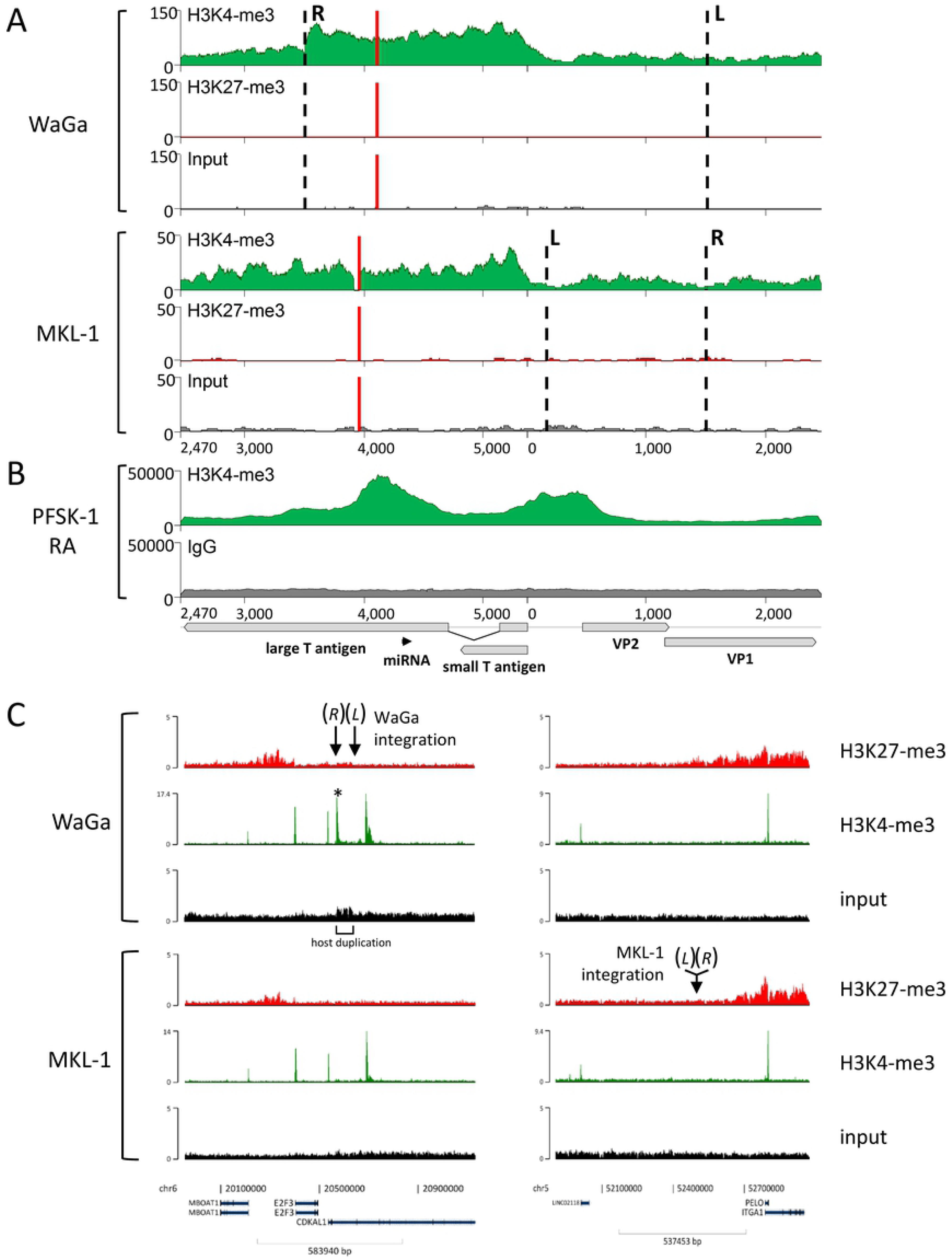
Histone modification pattern in MKL-1 and WaGa cells. (A): Coverage of the activating histone mark H3K4-me3 and the repressive histone mark H3K27-me3 on integrated MCPyV obtained by ChIP-Seq of WaGa and MKL-1 cells. (B): H3K4-me3 ChIP-Seq data from a replication assay (RA) performed in PFSK-1 cells were published before [37] and are included for comparison. Dashed lines represent breakpoints into the host genome, red lines the truncating event in LT. Note: The viral reference genome JN707599 is presented starting with nucleotide 2,470 for better visualization of ChIP-Seq patterns (see annotation of X-axis). (C): ChIP-Seq data for H3K4-me3 and H3K27-me3 from WaGa (upper panel) and MKL-1 cells (lower panel). The left and the right panel represent the two host genomic regions (1mbp) of the WaGa (left) and MKL-1 (right) integration sites. The corresponding junctions (*L* and *R*, marked by arrows) are indicated. The asterisk marks an additional H3K4-me3 signal which is not present in MKL-1. The signal is located within the 66kbp host duplication and flanks junction R. It originates from the H3K4-me3 signal of the early region of the integrated MCPyV genome that harbors the right breakpoint (R, see A) and extends into the host chromatin. Host duplication in WaGa is visible by the marked enhanced ChIP input signal.

Fig 8C shows H3K4-me3 and H3K27-me3 profiles across a 1mbp host region centered on the integration sites in WaGa and MKL-1 cells. Profiles for each site are shown for both cell lines to allow cross-comparison of epigenetic profiles. The overall profiles are almost identical, with the exception of WaGa cells showing an additional H3K4-me3 peak upstream of the integration site (marked with an asterisk in Fig 8C). The peak originates from H3K4-me3 signal at the right junction of the integrated MCPyV that spreads into the host. Further analysis of RNA-Seq data from WaGa and MKL-1 cells did not provide evidence for significant expression changes of CDKAL1 in WaGa compared to MKL-1 cells (S6C Fig). This result suggests that integration and establishment of additional intronic H3K4-me3 marks did not have immediate consequences for transcriptional regulation of the host gene.

### Epigenetic properties of MCC cell lines and MCPyV integration sites

We further sought to compare the global patterns of H3K4-me3 signals observed in WaGa and MKL-1 cells to other cellular entities, aiming to identify cell types, which may have a similar overall profile of this marks. Accordingly, we performed correlation and clustering analysis of the data from these two MCC cell lines in comparison to selected tumor cell lines and primary cells obtained from the ENCODE database. Our analysis revealed that both MCC cell lines show the highest correlation with each other. Next closest by hierarchical clustering are HeLa cells, mesenchymal stem cells, endothelial cells of the umbilical vein and fibroblasts of the dermis and lung (Fig 9A).

**Fig 9.**
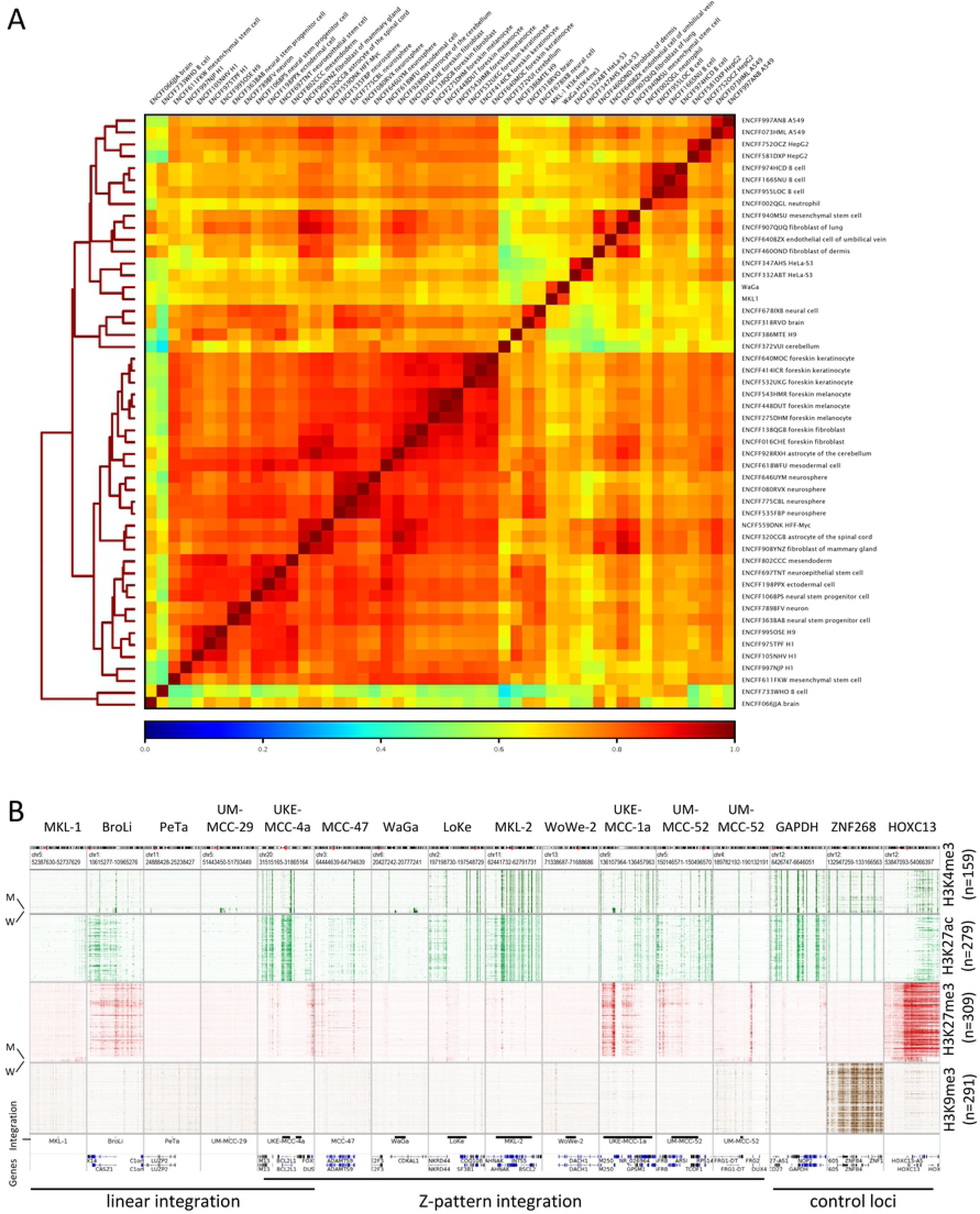
Epigenetic properties of MCC cell lines and MCPyV integration sites. (A): Correlation and clustering of H3K4-me3 profiles from WaGa and MKL-1 in comparison to 48 selected tumor cell lines and primary cells obtained from the ENCODE database. Correlation and clustering were performed using DeepTools and are based on MACS2 identified H3K4-me3 peak regions in the WaGa cell line. (B): Cellular chromatin environment at integration sites of MCC cell lines (350kbp window). Heat maps represent ENCODE ChIP-Seq signals of different cell types and cell lines (n is given beneath each modification) and include MKL-1 (M) and WaGa (W) data as indicated for H3K4-me3 and H3K27-me3 (please note increased track height of MKL-1 and WaGa for better visualization). Start and end of the bars in the integration track indicate positions of the left and right junctions of the respective integration site. Endogenous positive control regions were included for each histone modification using the same magnification (GAPDH: H3K4-me3 and H3K27-ac; ZNF268: H3K9-me3; HOXC13: H3K27-me3).

To investigate if MCPyV integration sites may possess general epigenetic features that may predispose them for integration we compared H3K4-me3, H3K27-ac, H3K27-me3 and H3K9-me3 profiles from selected cell lines and H3K4-me3 and H3K27-me3 profiles from MKL-1 and WaGa cells at the 13 MCPyV integration sites identified in our study (Fig 9B). We find that the integration loci are devoid of the histone modification H3K9-me3 (heterochromatin) in all cell lines, indicative of viral integration predominantly occurring in open chromatin structures. Similarly, most integration loci, except for BroLi, UKE-MCC-1a and UM-MCC-52, are devoid of the facultative heterochromatin mark H3K27-me3 in the majority of analyzed cell lines. The activating histone marks H3K27-ac and H3K4-me3 are present in seven out of 13 integration loci in most cell lines. The H3K27-me3 and H3K4-me3 profiles from WaGa and MKL-1 are in accordance with the majority of the other cell lines at most integration loci. These data suggest that integration of MCPyV favors open chromatin loci, which show histone-marks that are in general associated with active transcription. These features can be observed in the majority of cell lines analyzed by us including the MCC cell lines WaGa and MKL-1.

## Discussion

We here present a detailed analysis of MCPyV integration sites in 11 MCC cell lines and one primary tumor and its subsequent metastasis. Our study identifies two principal groups of integration patterns: (i) a linear integration of a single genome or viral genome concatemers and (ii) complex integration of single viral genomes or concatemers with duplications of adjacent host regions (Z-pattern), sometimes combined with additional rearrangements or amplifications. Within the linear integration groups, virus-host junctions are in close proximity (4 -18bp in the samples studied here), whereas more distant junctions (>17kbp in most cases) are typically observed for Z-pattern integrations.

In several cell lines, long contiguous nanopore sequencing reads (∼40-100 kbp) spanning complete integration loci provide direct evidence for the proposed integration site structure and permit exact determination of viral copy numbers of integrated concatemers. We further show that short-read capture sequencing allows distinction between integration of single viral genomes or concatemers, as well as accurate estimation of viral copy numbers.

We identified a single viral integration site in all samples but UM-MCC-52 and UKE-MCC-4a. These lines contain two integration sites each, but while viral sequences map to a single locus on Chr20 in UKE-MCC-4a, they are found on different chromosomes in UM-MCC-52. In UM-MCC-52, Chr5 contains a concatemeric integrate, whereas viral sequences integrated on Chr4 represent a partial genome that lacks the entire NCCR and 5’-proximal coding regions of the late and early genes. Since this partial genome contains three SNPs that are not found in concatemers on Chr5, it is likely that the two sites result from two independent integration events, with only the viral sequences on Chr5 contributing to transformation and tumorigenesis. Notably, we find that in all cases with viral concatemeric integrates, including the complex locus in sample UKE-MCC-4a, each viral genome copy carries identical, sample-specific LT-truncating mutations. This observation strongly supports the hypothesis that inactivating mutations occur prior to integration [20, 30], a model for which direct corroborating evidence as provided here has been missing thus far.

We furthermore propose that both LTtrunc mutations and viral concatemerization result from a similar mechanism as it has been previously suggested for papillomavirus integration [42] (Fig 10A). During the onset of viral DNA replication, polyoma- and papillomaviruses are thought to employ bidirectional theta replication to amplify their genomes. In this replication mode, two replication forks move in opposite direction along the episome, starting from the origin of replication in the viral NCCR. Normally, the replication complexes dissociate from viral DNA after collision of the two replication forks. However, if one of the forks stalls, the progressive fork may instead displace the 5’-end of the DNA synthesized by the stalled replication fork, resulting in a switch to rolling circle amplification (RCA). While herpesviruses encode factors (the viral terminase complex) which allow cleavage of unit length genomes from RCA products [43], such factors are missing in papilloma- and polyomaviruses. The missing cleavage activity therefore leads to the production of linear concatemers containing multiple viral genomes (with identical mutations) in a head-to-tail orientation. Previous reports on SV40 furthermore indicate that DNA replication stalls at preferred sites on the SV40 genome [44]. It is therefore conceivable that replication fork convergence leads to replication stalling within the early region, which thereby may represent a fragile site in which mutations, insertions and deletions occur resulting in truncated LT proteins. In addition to the linear concatemeric genomes with identical variants observed in the majority of MCC cell lines, we identified viral genomes with large inversions in the primary tumor MCC-47T, its descendent metastasis and the cell line PeTa (S5 Fig). Indeed, an *in vitro* SV40 replication model previously published by Ellen Fannings group [45] supports these observations. The study demonstrated that upon inhibition of the DNA repair protein ATM, SV40 replication favors RCA due to the continuous replication of one replication fork (Fig 10A, top). ATR inhibition, on the other hand, induces a dsDNA break when a moving replication fork collides with a stalled fork, resulting in broken replication intermediates (Fig 10A, bottom). Further recombination of such intermediates may lead to large inversions such as those that result in LT truncation in MCC-47T and PeTa (S5 Fig). Indeed, studies of MCPyV demonstrated ATM and ATR accumulation in viral replication centers and reported decreased viral DNA replication upon inhibition of these factors [46]. Hence, limitation of ATM and ATR during replicative stress might be responsible for the production of linear and defective MCPyV concatemers that we find integrated in the host DNA of MCCs.

**Fig 10.**
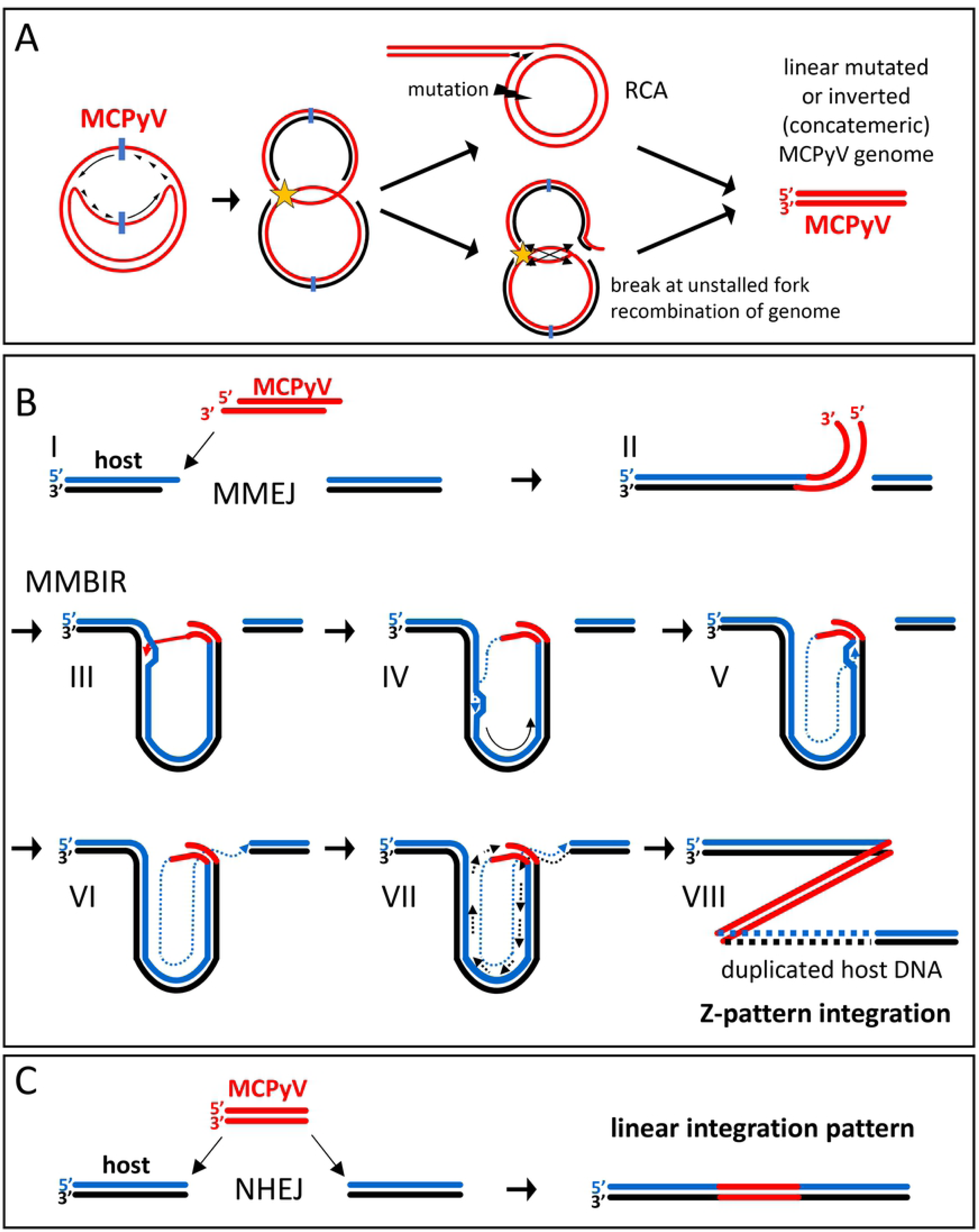
MCPyV integration model. (A): DNA replication of MCPyV is bidirectional (theta amplification) with replication forks starting at the ori (blue) and moving into opposite directions. Stalling replication forks (yellow star) can result in aberrant defective viral genomes. Top: Stalling replication forks induce mutations (black bolt) in the early region of the viral genome. The remaining fork induces unidirectional rolling circle amplification (RCA) resulting in large linear concatemers of mutated viral genomes. Bottom: Collision of a moving fork with a stalled fork leads to a dsDNA break at the moving fork. Recombination at the converging forks results in viral genomes with large inversions that truncate the early region. Both scenarios (RCA and break with recombination) yield linear defective (concatemeric) viral genomes. (B): (I) a linear viral genome is recognized as ds DNA break and undergoes resection of the 5’ ends by the host machinery. The same mechanism resects the 5’ end of a dsDNA break in the host DNA. (II) Homologies between viral and host sequences are used by microhomology-mediated end joining (MMEJ) to ligate the viral genome to a dsDNA break in the host genome. (III) The 3’ ss end of the viral genome invades a homologous host region and (IV) starts DNA synthesis in a D-loop structure (microhomology-mediated break-induced replication, MMBIR). (V) DNA synthesis reaches the original ds break with the viral genome and (VI) connects with the other side of the ds break by an unknown mechanism. (VII) The complementary strand is synthesized in a conservative mode using the newly synthesized strand as a template resulting in (VIII) an amplification of several kbp of host sequence surrounding the MCPyV integration site and a Z-pattern integration. (C): Without resection of 5’ ends a defective linear viral genome is integrated into a ds break of host DNA by nonhomologous end-joining (NHEJ). The integration mechanism is independent of homologies between viral and host sequences and results in a linear integration pattern.

While the above model provides a convenient explanation for mutagenesis and concatemerization of viral genomes, it does not explain how the integration process produces the distinct linear and Z-pattern integration patterns observed in our study. Motivated by studies of HPV integration sites in cervical cancer [39], we therefore searched for regions of microhomology between virus and host sequences at or nearby the virus-host junctions. We observed that in samples with a Z-pattern integration, the homology was significantly higher at the viral side of the junction compared to random sequences. This is not the case at the host side of the junction or in linear integrations in general. The lack of microhomologies on the virus side in linear integration also implies that the mechanisms leading to linear or Z-pattern integration already differ during the initial integration step. Based on our findings we propose a model in which two different pathways lead to the distinct MCPyV integration patterns observed in our study (Fig 10B and C). In both pathways, linear mutated single viral genomes, concatemeric genomes or recombined broken replication intermediates are the starting point (Fig 10A). We propose that Z-pattern integration begins with microhomology-mediated end joining (MMEJ) of a defective viral genome to a dsDNA break in the host DNA (Fig 10B). Therefor the viral DNA fragment is resected at the 5’ end, which distinguishes this pathway from nonhomologous end-joining (NHEJ) [47, 48]. The free 3’ end of the viral DNA aligns to a homologous region of a dsDNA break in the host genome which also underwent 5’ resection. The other end of the viral genome invades, again mediated by microhomologies, the host DNA upstream or downstream of the initial ds break. Subsequently, DNA synthesis starts using the host DNA as a template. This process is termed microhomology-mediated break-induced replication (MMBIR) and known to be involved in the amplification of large genomic regions (kbp to mbp range) in genetic disorders [31, 49, 50]. MMBIR has also been suggested as a mechanism in papillomavirus integration [39]. During MMBIR, the invading viral DNA strand is elongated in a so-called D-loop structure that uses the host DNA strand as a template as it moves forward. We hypothesize that DNA-synthesis proceeds until it reaches the position of the initial integration site of the viral DNA where two options exist. i) DNA synthesis continues using the viral DNA as a template leading to further amplification of host DNA together with viral DNA or ii) the nascent DNA strand connects to the other side of the original ds break and terminates the reaction. Currently, we can only speculate about the nature and accuracy of the mechanism mediating the ligation with the other side, as we obtained no reads covering these junctions. Nevertheless, our data from WaGa cells do not suggest a second round of amplification as we do not find appropriate copy numbers of viral and host DNA. To synthesize the complementary strand MMBIR then uses a conservative mode of DNA replication using the newly synthesized strand as a template [51, 52]. It is not clear if this occurs discontinuously by Okazaki fragments or continuously primed from the other side of the ds break [49]. Eventually, MMBIR results in the amplification of kbp of host DNA that we observe in Z-pattern integration. Furthermore, MMBIR could also explain the complex integration loci we observed for UM-MCC-52 and UKE-MCC-4a as frequent cycles of strand invasions with amplification of shorter stretches of DNA at each site have been reported for this mechanism [53]. S7 Fig shows the possible events that lead to the integration pattern in UM-MCC-52. Since we do not find significant microhomologies between viral and host sequences in the case of linear integration of MCPyV (Fig 10C), we propose that in this case viral sequences are integrated into a ds break of host DNA by NHEJ.

We also performed ChIP-Seq analysis of the histone modifications H3K4-me3 and H3K27-me3 in MKL-1 and WaGa cells. Interestingly, we find that the two cell lines exhibit strikingly similar modification profiles, suggesting that they share a distinct and MCC-specific epigenetic pattern. This notion is supported by hierarchical clustering analysis of H3K4-me3 profiles, which demonstrates very close relationship of MKL-1 and WaGa when compared to ENCODE datasets from 48 cell lines and primary cells. Of note, among the latter we find HPV positive HeLa cells being most similar to the MCC lines. Since MCPyV- and HPV-encoded oncoproteins interfere with similar cellular transformation, this may indicate that the epigenetic profile of WaGa and MKL-1 cells is being dominantly shaped by the viral oncoproteins. The next closest related H3K4-me3 profiles are from mesenchymal stem cells and fibroblasts of the lung and dermis, a finding which may support previous suggestions on the origin of the cells giving rise to MCC [54, 55]. We also find that neural cells cluster with the MCC cell lines, albeit more distantly than the cell lines and types mentioned above. Interestingly, a recent study reported reversion of MCC tumor cells to neuron-like cells after T-Antigen knockdown, suggesting neural precursor cells as putative MCC origin [56].

Our analysis of the epigenetic chromatin states at integration breakpoint positions in MCC cell lines and ENCODE datasets suggests that these regions are generally devoid of facultative or constitutive heterochromatin marks such as H3K27-me3 and H3K9-me3, but instead tend to carry euchromatin histone marks such as H3K4-me3. Similar to what has been reported for HPV [57, 58], our data thus suggest that MCPyV predominantly integrates into transcriptionally active regions characterized by open chromatin.

The ChIP-Seq analyses of activating and repressive histone marks in WaGa and MKL-1 cells further allowed us to investigate how gene expression from integrated viral genomes may be regulated. At least in these two cell lines, we do not find evidence for major alterations of overall host chromatin structure at the integration site. While we do not find repressive H3K27-me3 marks on integrated MCPyV genomes, we observe H3K4-me3 across the entire early region, a pattern which is markedly different from the more distinct peaks on the NCCR and the viral miRNA promoter of actively replicating episomes. Interestingly, a recent meta-analysis including >200 datasets from ChIP-Seq and ChIP-on-ChIP data found that broader H3K4-me3 peaks predict cell identity and are positively correlated with transcriptional consistency and precision [59]. While the exact molecular mechanisms which lead to formation of broad H3K4-me3 peaks are not yet clearly defined, we hypothesize that by this mechanism of buffering RNApol II pausing, stable viral oncoprotein expression is ensured.

In summary, we here report the first high-resolution analysis of MCPyV integration patterns in Merkel Cell Carcinoma using a combination of short- and long-read sequencing technologies. Our data strongly suggest a central role of microhomologies and DNA repair pathways including NHEJ and MMBIR in promoting either linear or Z-pattern integration. Our findings may not only explain MCPyV integration, but also substantiate previously suggested modes of human papillomavirus integration and viral-host DNA amplification mechanisms. Thus, our data suggest a common mechanism for papilloma- and polyomaviruses integration and provide the basis for further experimental studies to investigate the molecular events controlling this process.

## Methods

### Cell lines and tumor tissues

MCC cell lines WaGa [40], BroLi [40], LoKe [40], MKL-1 [60], MKL-2 [61], WoWe-2 [62], PeTa [62] were described before and were cultivated in RPMI 1640 with 10% FCS, 100 U/mL penicillin and 0.1 mg/mL streptomycin. For BroLi cells 20% FCS was used. UKE-MCC-1a and UKE-MCC4a were established in the department of dermatology at the Ruhr University of Essen. UM-MCC-29 and UM-MCC-52 have been described previously [63]. MCC-47 primary tumor and the corresponding bone metastasis tissue were isolated from an MCC patient (ID: 47) already described [64].

### DNA isolation

Genomic DNA used in capture sequencing was isolated applying the DNeasy Blood and Tissue Kit (Qiagen, Hilden, Germany) according to manufacturer’s instructions. For the isolation of HMW DNA used in nanochannel and nanopore sequencing cells were washed once in PBS, rotated for 10min at 4°C in nuclear extraction buffer 1 (50mM HEPES-KOH pH 7.5, 140mM NaCl, 1mM EDTA, 10% Glycerol, 0.5% Nonidet-P40, 0.25% Triton-X-100) and centrifuged for 5min at 2000xg. Pelleted cells were resuspended in nuclear extraction buffer 2 (10mM Tris-HCl pH 8.0, 200mM NaCl, 1mM EDTA, 0.5mM EGTA), rotated for 10min at 4°C followed by 5min at 2000xg. Pelleted nuclei were resuspended in nuclear extraction buffer 3 (10mM Tris-HCl pH 8.0, 10mM NaCl, 10mM EDTA, 1% SDS, 200µg/mL Proteinase K) and incubated overnight at 37°C. The mixture was subjected to two rounds of phenol extraction, one round of Phenol/Chloroform/Isoamyl alcohol (25:24:1) extraction and 2 rounds of chloroform washing steps. DNA was precipitated by adding 0.5 volumes of 2-propanol and 0.05 volumes of 5M NaCl solution. DNA was winded up using a small glass rod, washed in 70% Ethanol, air-dried and resuspended in TE-buffer (10mM Tris-HCl pH 8.0, 1mM EDTA).

### Capture sequencing

Capture probe sequencing of genomic DNA from MCC cell lines and tissues was performed using the SureSelectXT Target Enrichment System for Illumina Paired-End Multiplexed Sequencing Library (Agilent Technologies, Santa Clara, CA, USA) according to manufacturer’s instructions (protocol version B5). MCPyV-specific capturing probes were generated by shifting 120nt windows across the circular MCPyV genome (JN707599) with a step size of 1nt resulting in 5387 capture probes (S2 Table). Concentrations of all generated capture samples were measured with a Qubit 2.0 Fluorometer (Thermo Fisher Scientific, Waltham, Massachusetts, USA) and fragment lengths distribution of the final libraries was analyzed with the DNA High Sensitivity Chip on an Agilent 2100 Bioanalyzer (Agilent Technologies). All samples were normalized to 2nM and pooled equimolar. The library pool was sequenced on the MiSeq (Illumina, San Diego, CA, USA) with 2×150bp, with approximately 1.3mio reads per sample.

### Integration site identification from capture sequencing data

Illumina adapter sequences were removed from the capture sequencing short paired-end reads using cutadapt v2.7 [65]. Reads were then aligned to the MCPyV reference genome (JN707599) and the human reference genome (hg38) using minimap2 v2.14 [66] with the pre-set option --x sr. This option aligns short reads that may include indels or mismatches, e.g. LT antigen stop or frameshift mutations. Furthermore, the algorithm keeps unaligned parts of the reads and indicates these events as soft clipped bases within the SAM format CIGAR string. Furthermore, the option to report secondary alignments on virus and host was used to detect potential intra-viral fusions and rearrangements. After removal of PCR duplicates using samtools v1.9 [67], soft clipped reads, which indicate potential virus-host junctions or virus-virus rearrangements, were filtered by their respective CIGAR string. We set a minimum requirement of at least three independent unique reads containing the same potential breakpoint followed by the same consecutive soft clipped bases to detect putative virus-host junctions. The soft clipped portions of the reads were then again aligned to the human reference genome (hg38) using BLAST [68] and compared to the previous full length read alignments. The usage of those two different alignment methods resulted in high confidence junction sites (Table 1). Detected virus-host junctions were further confirmed by conventional PCR (S5 Table) and Sanger sequencing and/or Nanopore sequencing in selected samples. As described below (see variant detection) the cell lines MKL-1, LoKe, PeTa and WoWe-2 showed a contamination with WaGa DNA. For integration site detection in these cell lines all fusion reads originating from the contamination with WaGa were excluded.

### Variant detection

We used the aligned capture sequencing data to perform variant detection of viral integrates simultaneously in all samples with freebayes v1.3.1 [69], which reports the counts of reference as well as variant bases for each sample in vcf format. Frequencies were then calculated for each variant based on the count ratio of variant to total reads covering that position. The results are given in the S1 Table. Variants occurring with >99% frequency were counted as confidential variants. In four MCC cell lines three mutations identified as WaGa specific mutations (3,109 A to G, 3,923 G to A, 4,122 G to A) were detected in different frequencies (MKL-1: 9.5 to 18.6%; LoKe: 89% to 96.1%; PeTa: 78.1% to 96.1%; WoWe-2: 1.6% to 4.2%), which are indicative of a contamination prior capture hybridization. In these cell lines the WaGa specific variants were excluded and the cut-off for confidential variant detection was lowered according to the percentage of WaGa contamination.

### Nanopore sequencing

MinION sequencing libraries were generated from HMW DNA with the 1D genomic DNA by ligation kit (SQK-LSK109, ONT) according to the manufacturer’s instructions. MinION sequencing was performed as per the manufacturer’s guidelines using R9.4 flow cells (FLO-MIN106, ONT). MinION sequencing was controlled using Oxford Nanopore Technologies MinKNOW software. Base calling was performed using Guppy base calling Software v3.3.3 (ONT). Long reads were then aligned to the MCPyV reference genome (JN707599) as well as the human genome (hg38) using minimap2 [66] with pre-set parameters for MinION reads (-x map-ont). The S4 Table contains MinION sequencing details and summaries of quality, read numbers and sequence lengths.

### Nanochannel Optical Mapping

HMW DNA was labelled with two MCPyV specific probes (LT1: ATTO647N-GGCTCTCTG-CAAGCTTTTAGAGATTGCTCC; LT2: ATTO647N-GGCAACATCCCTCTGATGAAAGCTGCT-TTC) using the following components in an 80µl reaction: 4µg HMW DNA, 10µM of each random oligonucleotide pdN6, pdN8, pdN10, pdN12, pdN14, 16µl RT reaction buffer (5x), 0.5mM dNTPs, 1µM LT1 and LT2 each. The reaction was incubated for 10min at 95°C, followed by a decrease to 45°C in steps of 5°C per minute and 5min at 4°C. 4µl ReverTAid H Minus RT (Thermo Fisher Scientific) were added, incubated for 45min at 42°C and the reaction purified with 15µl of MagAttract beads (Qiagen). The labelled DNA was eluted with TE-buffer (10mM Tris-HCl pH 8.0, 1mM EDTA) and subsequently stained with a non-selective intercalating dye (TOTO-3 Iodide (642/660), Thermo Fisher Scientific) in a ratio of 1 dye every 5 bp to visualize the DNA fragments by fluorescence microscopy. The nanofluidic devices for the measurement were made by direct imprinting in Ormostamp (a commercial, UV-curable polymer, micro resist technology GmbH, Berlin, Germany) as explained elsewhere [70-72]. They contain two U-shaped microchannels to deliver the molecules from the inlets into the nanochannels and 3D-tapered inlets to connect the micro and nanostructures, pre-stretch the molecules and avoid clogging. The nanochannels are 280 nm wide, which is in the order of the DNA persistence length (50 nm); the molecules are elongated and significantly stretched, (∼25 % of their full contour-length in this particular case). The flow of the molecules is observed in an inverted, fluorescent microscope (TiU, Nikon, Tokyo, Japan) using an EM-CCD Camera (Evolve Delta, Photometrics, Tucson, AZ, USA) with a 100x oil immersion objective. The real-time signal is obtained using a laser beam (λ=633nm, 0.2mW excitation power) focused on the central part of the nanochannel by the objective. The emitted fluorescence signal is recorded in real-time with a single photon counter (COUNT Module, Laser Components GmbH, Olching, Germany), while the excitation signal is filtered out by using a spectral filter (692/40nm band-pass filter, Semrock, Rochester, NY, USA). In this configuration, the molecules are detected as step-like peaks in time scans, allowing for real-time read-out with high throughput. Peak analysis (as photoluminescence intensity and duration time) gives information about the molecule length, as well as its genome-dependent barcode.

### ChIP-Seq analysis

ChIP assays were performed as previously described [37, 73] with the following changes. For each IP 100µL chromatin was pre-cleared with BSA blocked protein-G sepharose beads (GE Healthcare, Chicago, IL, USA) and incubated for 16h at 4°C with 2µL α-H3K27me3 antibody (#07-449; Merck Millipore, Burlington, MA, USA) or α-H3K4-me3 antibody (Rabbit monoclonal antibody (#04–745, clone MC315; Merck Millipore). DNA was purified by phenol-chloroform extraction and ethanol precipitation. ChIP and corresponding input libraries were prepared from 2–10 ng DNA using the NEXTflex Illumina ChIP-Seq Library Prep Kit (#5143–02; Bioo Scientific, Austin, TX, USA) according to the manufacturer’s instructions. Illumina libraries were sequenced on a NextSeq 500 (Illumina) using single-read (1×75) flow cells at a sequencing depth of 30Mio reads.

Quality filtered single end reads were aligned to the viral reference genomes of MCPyV (JN707599) and human genomes (hg38) using Bowtie [74] with standard settings. Coverage calculation for visualization purposes was performed with IGV-Tools [75]. Visualization was performed using IGV and EaSeq [76].

### RNA-Seq analysis

RNA-Seq analysis of WaGa and MKL-1 cells was performed essentially as described previously [24]. Briefly, high quality RNA of both cell lines was subjected to Illumina compatible library preparation. Libraries were then sequenced on an Illumina HiSeq2500 and analyzed using STAR splice aware read mapping. DEseq2 was used to perform differential gene expression analysis.

### Chromosome copy number variation analysis

Genome-wide chromosome copy number variation analysis in MKL-1 and WaGa cells was performed with FREEC [77] using low coverage WGS data (ChIP-input). Sequencing data of female HDF cells were used as normal chromosome set control (2n). Counting windows during FREEC analysis was set to 50.000 bp. Visualization of genome-wide copy number variation data was then performed with circos [78]. Color-coding of the shown circos plot indicates 2n (black), 1n (green) and >= 3n (red) chromosomal regions.

### Analysis of host region amplification

To calculate, how often the host region within the two WaGa specific integration sites is duplicated and whether host regions preceding or following the integration site in MKL-1 exhibit differential copy numbers, regions of 60kb in length (size of the host duplication in WaGa) were divided into overlapping regions of 5kb with a shift size of 2.5kb. Reads from WaGa and MKL-1 low coverage WGS samples (ChIP input) were counted using featureCounts [79]. All length normalized counting data were then normalized within each sample to the median read counts of the measured region of Chr3, which represents 2n in both cell lines according to copy number variation analysis described above.

To estimate the copy number of the host amplifications in UKE-MCC-4a we counted MinION reads (> 3 kbp) covering the host integration locus (R I to L I) on chr20 and compared it to random control loci of the same size (120 kbp) on chromosomes 3, 4 and 5 using featureCounts. The number of selected control regions varied between 12 and 40 sites per chromosome according to the respective chromosome size. For comparison, the median value of chr3 control loci was set to 1.

### Calculation of concatemeric unit counts

Numbers of MCPyV concatemers in WaGa, MKL-1, UKE-MCC-4a and UM-MCC-52 were derived directly from nanopore sequencing reads spanning the entire integration. Besides, for MKL-1 and WaGa, concatemer count numbers were calculated from low coverage WGS data (ChIP Input). The viral reference genome was divided into overlapping 1kbp windows with a shift size of 0.5kbp and reads were counted using featureCounts [79]. Length normalized counts were then additionally normalized to both, the count data of Chr3 as well as the total number of integrations per cell (WaGa = 2 due to the host duplication of Chr6; MKL1 = 1).

To estimate the viral genome numbers in concatemers of MCPyV integrates in the remaining samples we counted the capture sequencing reads containing the 25 virus-specific bases at each virus-host junction indicating the virus coverage (i.e. virus only plus fusion) next to the junction (A). Additionally, we counted all junction-spanning reads containing 22 virus-specific followed by 3 host-specific bases (B); note: this imbalance of viral and host bases is necessary to avoid capture sequencing-introduced bias. The estimated number of concatemeric full-length units (F) from each breakpoint can be calculated as F = ||A/(A-B)-1||. This formula is restricted to samples with more than one full-length unit and with breakpoints separated by at least 25 bases. For each MCC sample, we presented the range of estimation results of all detected breakpoints (Table 1).

### Correlation and cluster analysis of ChIP-Seq and ENCODE data

All ENCODE [80] dataset information used in this study is given in the S3 Table. We remapped the reads from WaGa and MKL-1 input and H3K4-me3 samples to maximize comparability with the ENCODE data using the ENCODE ChIP-Seq-pipeline2 (https://github.com/ENCODE-DCC/chip-seq-pipeline2). The resulting pval bigWig data were then used for downstream analysis. For the subsequent analysis H3K4-me3 enriched sites were detected in WaGa and MKL-1 using MACS2 peak calling [81]. We performed Person correlation and cluster analysis with 48 selected H3K4-me3 ENCODE data sets (S3 Table) together with WaGa and MKL-1 using DeepTools v3.1.3 (multiBigwigSummary and plotCorrelation) [82]. Analysis was restricted to the WaGa H3K4-me3 peak regions to reduce the influence of background signal.

### Microhomology analysis

Sequences upstream and downstream of breakpoints were selected for microhomology analysis. For each breakpoint two sequences of 40bp, one of viral (seqVir) and one of human (seqHum) origin, were obtained. Each seqVir was compared with the sequence of the corresponding region in the human reference assembly (GRCh38). If a seqVir was observed upstream of the breakpoint and a seqHum was downstream of the breakpoint, the seqVir was compared to the sequence 40bp upstream of the seqHum in the human reference assembly (and vice versa). Likewise, each seqHum was compared with sequences from the viral reference assembly (JN707599). Next all 3-mers co-occuring in both sequences were identified. Paths connecting these k-mers were constructed maintaining the observed order of k-mers in both sequences. Paths containing not at least one pair of overlapping kmers were ignored. For each remaining path a score was calculated: score = 2 x bases_in_kmers – | positionviral + positionhuman| // 2. Positionviral and positionhuman are the positions (0-based) of bases which are located in kmers and which are closest to the breakpoint. Only the highest scoring paths were kept. Thus, two scores for each side of a junction were obtained, one for the comparison of seqHum with the viral reference sequence (scoreHum) and one for the comparison of seqVir with the human reference sequence (scoreVir). Unpaired two-tailed t-tests were applied for comparing the scoresVir and scoresHum of a selected group (linear or Z-pattern integration) with the scores obtained for 200 randomly selected genomic positions of the viral and the human reference sequence. To account for its circularity, the viral reference sequence was correspondingly prefixed and suffixed with 40bp before random selection. We excluded UKE-MCC-4a from the analysis due to its complex integration pattern, UM-MCC-52 was categorized as Z-pattern integration.

### Data availability

Sequencing data are accessible in the public repository ENA, accession number PRJEB36884.

## Acknowledgement

We are grateful to Kerstin Reumann, Marion Ziegler and Christina Herrde for excellent technical support.

## Conflict of interest

The authors state no conflict of interest.

## Supporting information

**S1 Fig. Virus-host junctions from integration sites of all samples with microhomologies between virus and host sequences.** Sequences at the virus-host junction (in grey) were derived from capture sequencing and aligned to reference sequences for the human genome (hg38) and MCPyV (JN707599). L=left side of the integrated viral genome, R= right side. Depicted are 40 bps upstream and downstream from the virus-host junction (indicated by a black line). In the case of insertions at the junction, sequences were extended for the length of the insertion. Human sequences are depicted in blue and viral sequences in black letters. Detected microhomologies (see material and methods) are marked in red.

**S2 Fig. Reads from capture sequencing of sample UM-MCC-52 are aligned to the MCPyV genome (JN707599).** Grey color represents perfect matching of read and reference sequence. Blue, red, green and orange show mutations in the read sequence to the bases C, T, A and G respectively. Breakpoints into the host genome are indicated at the top reflected by longer stretches of mismatching bases. Lower panels show magnification of alignment. Mutations at bp 1,792 and 1,816 (G to C, left panel, red arrows) are not present in reads leading into Chr5. Reads that contain these mutations also contain a G to C transition at bp 1,708 (green arrow). Mutations in LT including the inactivating mutation (stop) are present in all captured sequences (right panel).

**S3 Fig. Reads derived from capture sequencing of sample UKE-MCC-4a are aligned to the MCPyV genome (JN707599).** Color code is identical to S2 Fig. Breakpoints into the host genome are indicated at the top and can be recognized by longer stretches of mismatching bases. Bp 2,053 to 3,047 are deleted in approximately one third of the reads covering the region. This region also contains a breakpoint into the host genome indicating an integration of two versions of MCPyV (one with and one without a deletion). Mutations in LT including the inactivating mutation (stop) are present in all captured sequences.

**S4 Fig. Coverage profiles of the of the cell lines LoKe, PeTa, WoWe-2, UKE-MCC-1a, UM-MCC-29 and MCC-47T/M.** MCPyV-host fusion reads from capture sequencing were mapped to the human genome. (A): PeTa and UM-MCC-29 show a coverage profile characteristic for a linear integration pattern. (B): LoKe, WoWe-2 and UKE-MCC-1a show a coverage profile characteristic for a Z-pattern integration. (C): The sample MCC-47 (tumor and metastasis) shows a coverage profile with short distance (4bp) of breakpoints on the host genome but outward-facing orientation of viral sequences. The result is a Z-pattern integration with duplication of 6bp of host DNA as depicted in the right panel. Reads for both junctions of the tumor and the left junction of the metastasis are mapped by BLAST only.

**S5 Fig. Rearranged MCPyV genome and integration locus of sample MCC-47T/M.** (A): Rearranged MCPyV genome derived from capture sequencing of sample MCC-47 (primary tumor and metastasis) compared to MCPyV wild type (JN707599). For better comparison, both genomes are depicted as episomes. Breakpoints into the host genome are indicated (bp 5,193 and 5,290). Bp 1547-4119 are inverted with 1,547 fused to 4,166 and 4,119 to 991 causing a frameshift in LT that leads to a stop at position 4,166. The C-terminal part of LT fused to VP2 is also out of frame, which causes a stop at the beginning of the LT C-terminus. (B): Integration locus of MCC-47 derived from capture sequencing (chr3: 64,619,639-44). The rearranged MCPyV genome is integrated as a concatemer with at least one complete viral genome being flanked by partial genomes that connect into the host genome. 6bp of host sequence are duplicated at the integration site.

**S6 Fig. RNA-Seq analysis of WaGa and MKL-1 integration locus.** (A): Counts of host, virus and host-virus-fusion splice junction reads connecting to the splice acceptor of the second LT exon in MKL-1 and WaGa cells. In WaGa cells, we additionally counted splices between exons 4 and 5 of CDKAL1 (splice 3). It is likely that the transcripts harboring these splices originate from the copy of chr6 that does not contain the viral integrate. All detected splice events use annotated donor and acceptor sites as indicated. (B): RNA-Seq coverage at the integration locus. Detected splices are indicated by arcs. For further details, see legend to Figure 4. (C): Normalized RNA-Seq data of CDKAL1. Three RNA-Seq datasets of WaGa and MKL-1 (one dataset generated in this study and two datasets previously published [24]) were combined and subjected to standard DEseq2 analysis. Shown are Deseq2 normalized counts of CDKAL1 (n=3, mean + SEM). The slight Log2 fold change of 0.15 between both cell lines was found to be not significant (ns) by DEseq2 analysis.

**S7 Fig. Model for MCPyV integration in the complex integration locus of UM-MCC-52 on Chr5**. (I) Mutated concatemeric MCPyV genomes (at least 4 copies of MCPyV in this case) are produced by RCA and undergo 5’ resection by the host machinery. (II) Ligation to a ds break in the host DNA at the left side (L) is achieved by MMEJ. (III) The viral genome loops back and invades with its 3’ end a homologous host region and (IV) starts DNA synthesis in a D-loop structure (MMBIR). Different to the general model, the 3’ end of the viral genome aligns to the forward not the reverse strand. (V) DNA synthesis continues until it reaches site a and the D-loop disassembles. (VI) The newly synthesized strand invades again the host DNA (site b), this time aligning to the reverse strand. (VII) DNA replication can now proceed until it reaches L were it connects to the original ds break by an unknown mechanism (VIII). (IX) The complementary strand is synthesized in a conservative mode using the newly synthesized strand as a template. (X) For UM-MCC-52 the result is an amplification of several kbp of host sequence between L and b as well as an inverted sequence between site a and R.

**S1 Table. Variants in MCC sample derived MCPyV sequences compared to reference JN707599 obtained by capture sequencing.** The upper panel contains larger rearrangements and deletions. The lower panel contains small indels as well as SNPs. Blue fields indicate variants that occur in >99% of reads. Orange fields contain variants present in a subset of reads.

**S2 Table. Capture probes for MCPyV used in capture sequencing of MCC samples.**

**S3 Table. ENCODE data sets included in Figure 9.**

**S4 Table. Statistics of nanopore sequencing.**

**S5 Table. Primers used for sanger sequencing.**

